# Metabolic activities are selective modulators for individual segmentation clock processes

**DOI:** 10.1101/2024.06.04.597451

**Authors:** Mitsuhiro Matsuda, Jorge Lázaro, Miki Ebisuya

**Affiliations:** European Molecular Biology Laboratory (EMBL) Barcelona, 08003 Barcelona, Spain; Cluster of Excellence Physics of Life, TU Dresden, Dresden, Germany; Collaboration for joint PhD degree between EMBL and Heidelberg University, Faculty of Biosciences, Heidelberg, Germany; Max Planck Institute of Molecular Cell Biology and Genetics, Dresden, Germany

## Abstract

A sequence of cellular and molecular processes unfolding during embryonic development prompts fundamental questions of how the tempo of multiple processes is coordinated and whether a common global modulator exists. The oscillation of the segmentation clock is a well-studied model of developmental tempo. While the clock period is known to scale with the kinetics of gene expression and degradation processes of the core clock gene Hes7 across mammalian species, how these key molecular processes are coordinated remains unclear. In this study, we investigated if metabolic activities act as a global modulator for the segmentation clock, finding that they are rather selective modulators. While several metabolic inhibitions extended the clock period, their effects on the key processes varied. Inhibition of glycolysis decelerated the protein degradation of Hes7 and extended the production delay but did not influence the intron delay. Electron transport chain inhibition extended Hes7 intron delay without influencing the other two processes. Combinations of distinct metabolic inhibitions exhibited synergistic effects. By contrast, temperature changes affected the clock period and all three key processes simultaneously. These results highlight the selective effects of metabolic activities on segmentation clock processes, hinting that their scaled kinetics across species may be achieved through combinations of multiple modulators.

## Introduction

During the progression of embryonic development, multiple cellular and molecular processes happen sequentially and concurrently: while cells undergo proliferation, differentiation, and intercellular communication, a myriad of gene products are synthesized, processed, transported, and degraded in each cell. Since altering the timing or duration in one process without adjusting that in the others can perturb the precise spatiotemporal patterning of cells and tissues, coordinating tempo across multiple processes is crucial. This prompts the question of whether a common global factor exists to modulate the tempo of these processes simultaneously. Moreover, given that developmental tempo varies across animal species, another fundamental question is whether different species utilize the same global modulator.

There has been a debate on the existence and nature of a global modulator of developmental tempo^1–3^. For example, temperature can be a global modulator for ectotherms, such as insects, fish, and frogs^4–9^. As most biochemical reactions accelerate or decelerate by 2-3 folds for every 10-degree change in temperature, the external temperature can synchronously set the tempo of most cellular and molecular processes. By contrast, endotherms including mammals develop at relatively constant, similar temperatures, and thus these animals are unlikely to use temperature as a global modulator. Another attractive candidate for a global modulator is metabolism or energy. As all biological processes require energy, alteration in metabolism may influence multiple processes simultaneously. Indeed, mitochondrial activity is associated with several molecular and cellular processes, including gene expression, protein degradation, and neuronal development^10–13^. At the organismal level, nutrition is well-known to influence growth, and allometric metabolic rates are proposed to underlie diverse growth rates across species^14,15^. Mutants for mitochondrial components also display sluggish development in C. elegans and mice^16–19^. However, it remains unclear to what extent metabolism can modulate the tempo of multiple processes simultaneously.

The segmentation clock is widely used as a model to study developmental tempo. The oscillatory gene expression of the segmentation clock regulates the timings of periodic body segment formation during embryogenesis^20^. The segmentation clock has been recapitulated from pluripotent stem cells of multiple species by several groups, and the oscillation periods are reproducible despite variations in the differentiation protocols and cell lines^21–28^. The in vitro segmentation clock is amenable to detailed measurements of the kinetics of molecular processes. The core molecular mechanism of segmentation clock oscillation is a delayed negative feedback loop of the Hes7 gene (Fig. 1a): the Hes7 protein produced through gene expression processes represses its own promoter, before being degraded eventually. The negative feedback loop with delays can give rise to oscillatory gene expression of Hes7, and its oscillation period is mainly determined by the degradation rates and delays in the loop^29–31^; slower degradation and longer delays lead to a longer period. Even though the individual oscillations are further synchronized among cells, this study focuses on the cell-autonomous oscillation mechanism. A previous study demonstrated that the protein/mRNA degradation rates, the production delay (i.e., the delay derived from production processes such as transcription and translation), and the intron delay (i.e., the delay associated with intron sequences such as splicing) of the Hes7 gene could largely account for the cell-autonomous periods of mouse and human segmentation clocks (Fig. 1a)^25^.

**Figure 1.**
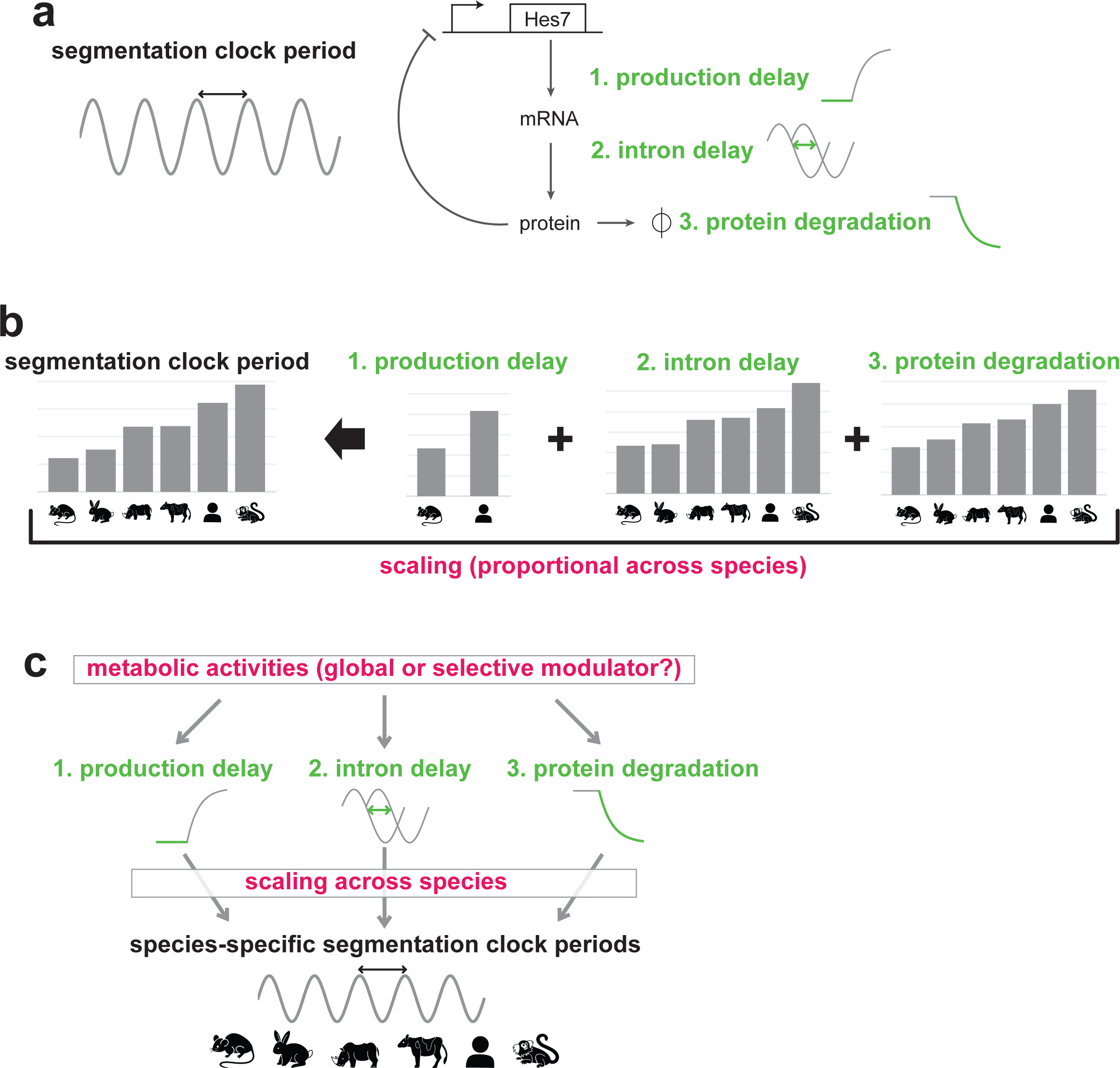
Scaling of the segmentation clock and the question. **a,** Schematic representation of the core mechanism of segmentation clock oscillation: a delayed negative feedback loop of the Hes7 gene. Hes7 is a transcriptional repressor that inhibits its own promoter, giving rise to the oscillatory expression. The production process of the Hes7 protein takes time due to transcription and translation (1. production delay) whereas the Hes7 intron processing also takes time (2. intron delay). The Hes7 protein is eventually degraded through the ubiquitin-proteasome pathway (3. protein degradation). The oscillation period, namely the segmentation clock period, is largely determined by the degradation rates of Hes7 mRNA/protein as well as the total delay in Hes7 gene expression (the sum of the production delay and intron delay). Hes7 mRNA degradation rate is not assessed in this study. **b,** The key kinetic parameters of Hes7 (1. production delay, 2. intron delay, and 3. protein degradation rate) are highly correlated with the segmentation clock period across mouse, rabbit, rhinoceros, bovine, human, and marmoset cells. This proportional change in the parameters is termed scaling in this study. Data are from Matsuda et al. (2020)^25^ and Lázaro et al. (2023)^27^. The production delay data are available only for mouse and human. **c,** This study addresses whether metabolic activities act as a global modulator that simultaneously affects the three key molecular processes of the segmentation clock or selective modulators.

The segmentation clock periods are notably species-specific (Fig. 1b); even among mammals, the periods are 2-3 h in mice and rabbits, ∼4 h in rhinoceroses, pigs, and cattle, ∼5 h in humans, and ∼6 h in common marmosets^20,22,24,25,27,28^. The three key kinetic parameters of the Hes7 gene, namely the protein degradation rate, intron delay, and production delay, are highly correlated with the segmentation clock periods across species (Fig. 1b)^25,27^. Although it is theoretically possible, for example, to extend the period by extending only the intron delay without altering the other two kinetic parameters, each species simultaneously and proportionally modulates the three parameters to give rise to the species-specific period. The remarkable scaling of these parameters implies but does not prove, the existence of a common global modulator that simultaneously controls the protein degradation, intron processing, and production processes of Hes7 across species. Alternatively, these individual molecular processes may have been modulated by separate mechanisms through evolutionary selection (Fig. 1c).

Metabolism is known to affect the segmentation clock. Modulation of glycolytic activity disturbs somitogenesis in embryos^32–36^ whereas modulation of electron transport chain (ETC) activity can alter the period of the in vitro segmentation clock^26,27^. These results raise a hypothesis that metabolism may be a global modulator that simultaneously controls the multiple molecular processes of Hes7 to tune the segmentation clock period (Fig. 1c). It must be noted, however, that metabolism is an umbrella term representing a broad spectrum of activities and phenomena, including ATP production, ETC activity, glycolysis, mTOR pathway signaling, metabolites, protein/RNA turnover, and even mitochondrial morphology. Indeed, recent studies reported that rather than ATP production, more specific metabolic factors, such as the cytosolic NAD+/NADH ratio, intracellular pH, and glycolytic metabolites, are crucial for the segmentation clock and somitogenesis^26,34–36^. As the metabolism hypothesis originally gained popularity because of the universal role of energy and ATP in biological processes, it is debatable if a specific metabolic activity can act as a global modulator for the key molecular processes of the segmentation clock (Fig. 1c).

In this study, we imposed different types of metabolic inhibitions on the segmentation clock. Unlike temperature change, the effects of metabolic inhibitions on the three key molecular processes of Hes7 were selective, casting doubt on the concept of metabolic activities as a global modulator for the segmentation clock tempo.

## Results

### Glycolysis or ETC inhibition selectively affects individual segmentation clock processes

We examined the effects of different types of metabolic inhibitors on the mouse segmentation clock period (Fig. 2a). An ETC inhibitor, sodium azide (azide) is reported to decelerate the segmentation clock^26,27^, and a glycolysis inhibitor, 2-Deoxy-D-glucose (2DG) is reported to perturb somitogenesis^33,34^. Mouse and human presomitic mesoderm (PSM) cells were induced from mouse Epiblast stem cells (EpiSCs) and human induced pluripotent stem cells (iPSCs), respectively, and the segmentation clock oscillation in PSM cells was visualized with the Hes7 promoter-luciferase reporter^23,25,27^. Glycolysis inhibition or ETC inhibition extended the mouse segmentation clock period in a dose-dependent manner (Fig. 2b,c). The periods in mouse PSM cells treated with 10 mM 2DG and 1 mM azide were 191 ± 2.1 min (mean ± sd) and 204 ± 8.8 min, respectively. While they were significantly longer than the control mouse period (152 ± 3.5 min), they were still much shorter than the human period (325 ± 5.5 min).

**Figure 2.**
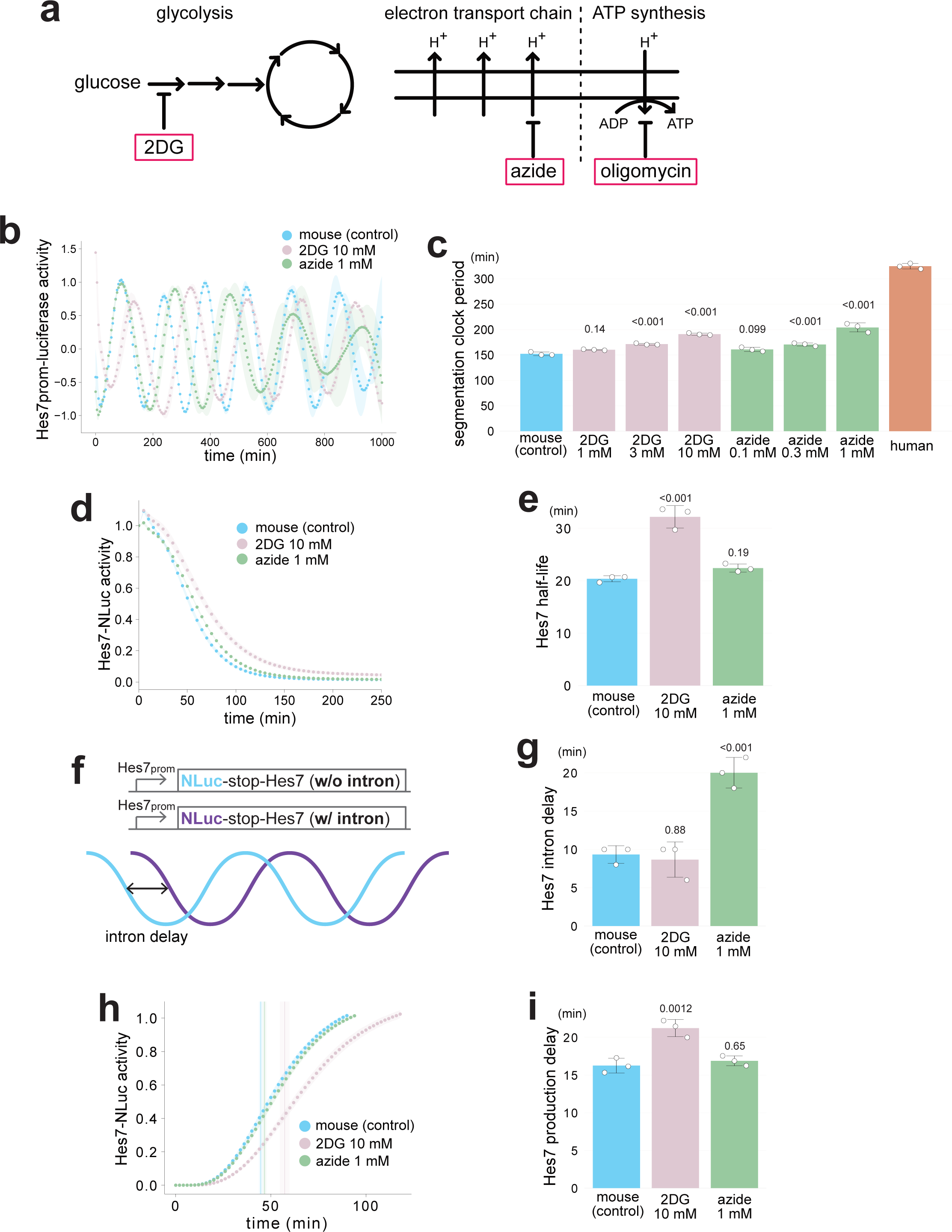
Selective effects of metabolic inhibitions on the key molecular processes of the segmentation clock. **a,** Schematic representation of cellular metabolic pathways and their pharmacological inhibitors. 2-Deoxy-D-glucose (2DG), sodium azide (azide), and oligomycin are known inhibitors of glycolysis, electron transport chain (ETC), and ATP synthesis, respectively. **b,** Dose-dependent effects of 2DG or azide on the segmentation clock period. Mouse PSM cells were pre-treated with the metabolic inhibitors for 2 h, and the oscillatory activity of the Hes7 promoter-luciferase reporter was monitored. The signal was detrended and amplitude-normalized. **c,** Hes7 oscillation periods estimated from b. Human data are from Fig. 5. **d,** Effects of 2DG or azide on Hes7 protein degradation. Mouse PSM cells were pre-treated with the inhibitors for 4 h. The transcription of the Hes7 reporter fused with Nano luciferase (NLuc) was halted upon the addition of doxycycline (Dox) at time 0, and the decay of the Hes7-NLuc signal was monitored. **e,** Hes7 half-lives estimated from d and Supplementary Fig. 1. **f,** Hes7 intron delay assay. The oscillation phase difference between the luciferase reporter without (w/o) Hes7 intron sequences and the one with (w/) Hes7 intron sequences was defined as the intron delay. **g,** Effects of 2DG or azide on Hes7 intron delay. Mouse PSM cells were pre-treated with the inhibitors for 4 h, and the oscillatory activities of the luciferase reporters w/o and w/ Hes7 intron sequences were monitored. Intron delays were estimated from Supplementary Fig. 2. **h,** Effects of 2DG or azide on Hes7 production delay. Mouse PSM cells were pre-treated with the inhibitors for 4 h. The transcription of Hes7-NLuc was induced upon Dox addition, and the onset of the Hes7-NLuc signal was monitored. Vertical lines indicate the inflection points. **i,** Hes7 production delays estimated from h and Supplementary Fig. 3. **b,d,h,** Shading indicates mean ± sd (n = 3). **c,e,g,i,** Graphs indicate mean ± sd (n = 3). P-values are from two-sided Dunnett’s test against the indicated controls.

We then examined the effects of metabolic inhibitions on the three key kinetic parameters of the Hes7 gene that mostly accounted for the segmentation clock period: the Hes7 protein degradation rate, intron delay, and production delay (Fig. 1b)^25^. Protein degradation was measured by halting the transcription of the Hes7-Nanoluciferase (NLuc) reporter and monitoring the decay in the luciferase signal (Supplementary Fig. 1a)^25^. Glycolysis inhibition decelerated Hes7 protein degradation (Fig. 2d,e; Supplementary Fig. 1b); the Hes7 half-life measured in the 2DG-treated mouse PSM cells was 32 ± 2.1 min while that in untreated cells was 20 ± 0.6 min. By contrast, ETC inhibition by azide treatment did not show a significant influence on Hes7 protein degradation (Fig. 2d,e; Supplementary Fig. 1b). The intron delay represents a combined delay caused by the Hes7 intron sequence, including the delay due to intron splicing. The intron delay was defined as the oscillation phase difference between two Hes7 promoter-luciferase reporters with and without Hes7 intron sequences; the oscillation phase of the reporter without introns should always precede the one with introns (Fig. 2f; Supplementary Fig. 2a)^25^. Azide treatment notably extended Hes7 intron delay; 20 ± 2.0 min in azide-treated mouse PSM cells as compared to 9 ± 1.2 min in untreated cells (Fig. 2g; Supplementary Fig. 2b). By contrast, 2DG treatment did not show a significant influence on Hes7 intron delay. The production delay represents a combined delay caused by the gene expression steps of Hes7, including transcription and translation, except for intron-related steps. The production delay was measured by inducing the expression of the Hes7-NLuc reporter and monitoring the onset timing^25^ (Supplementary Fig. 3a). While 2DG treatment extended Hes7 production delay (21 ± 1.1 min in 2DG-treated mouse PSM cells as compared to 16 ± 1.0 min in untreated cells), azide treatment did not show a significant influence (Fig. 2h,i; Supplementary Fig. 3b). These results demonstrated selective and disproportionate effects of glycolysis and ETC inhibitions on the three key molecular processes of the mouse segmentation clock even though the two metabolic inhibitors extended the period to a similar extent; glycolysis inhibition selectively affected Hes7 protein degradation and production delay whereas ETC inhibition affected only Hes7 intron delay.

### Glycolysis, ETC, and ATP synthesis inhibitions synergistically extend the clock period

The selective and complementary effects of glycolysis and ETC inhibitions prompted us to investigate whether their combined inhibition could yield synergistic effects on the segmentation clock period. While individual treatments with either 2DG or azide led to only a modest extension of the period, their combinations notably extended it (Fig. 3a-d). For example, the period of mouse PSM cells treated with 1 mM azide and 2 mM 2DG was 287 ± 13 min, resembling the human period more closely than the mouse (Fig. 3b). Even at lower concentrations (0.3 mM azide and 1 mM 2DG), each of which individually showed only negligible effects (Fig. 2c), the combined treatment resulted in a clearly extended period of 206 ± 0.7 min (Fig. 3d). It must be noted, however, that the combined metabolic inhibitions also lowered the reporter signal (Supplementary Fig. 4). These observations led us to hypothesize that even a metabolic inhibitor previously deemed ineffective on the segmentation clock might exhibit a notable impact when combined with another inhibitor. Oligomycin is an ATP synthase inhibitor (Fig. 2a), and it was reported to have no significant influence on the segmentation clock period^26^. As high concentrations of oligomycin cause the dampening of segmentation clock oscillation^26^, we employed low concentrations (10-100 nM). While oligomycin treatment alone exhibited a significant yet very modest extension of the period (e.g., 177 ± 1.5 min in mouse PSM cell treated with 10 nM oligomycin), combinations of oligomycin and 2DG notably extended it (e.g., 248 ± 2.9 min in mouse PSM cells treated with 1 mM 2DG + 10 nM oligomycin) (Fig. 3f). These results indicated synergistic effects of different types of metabolic inhibitions on the segmentation clock period.

**Figure 3.**
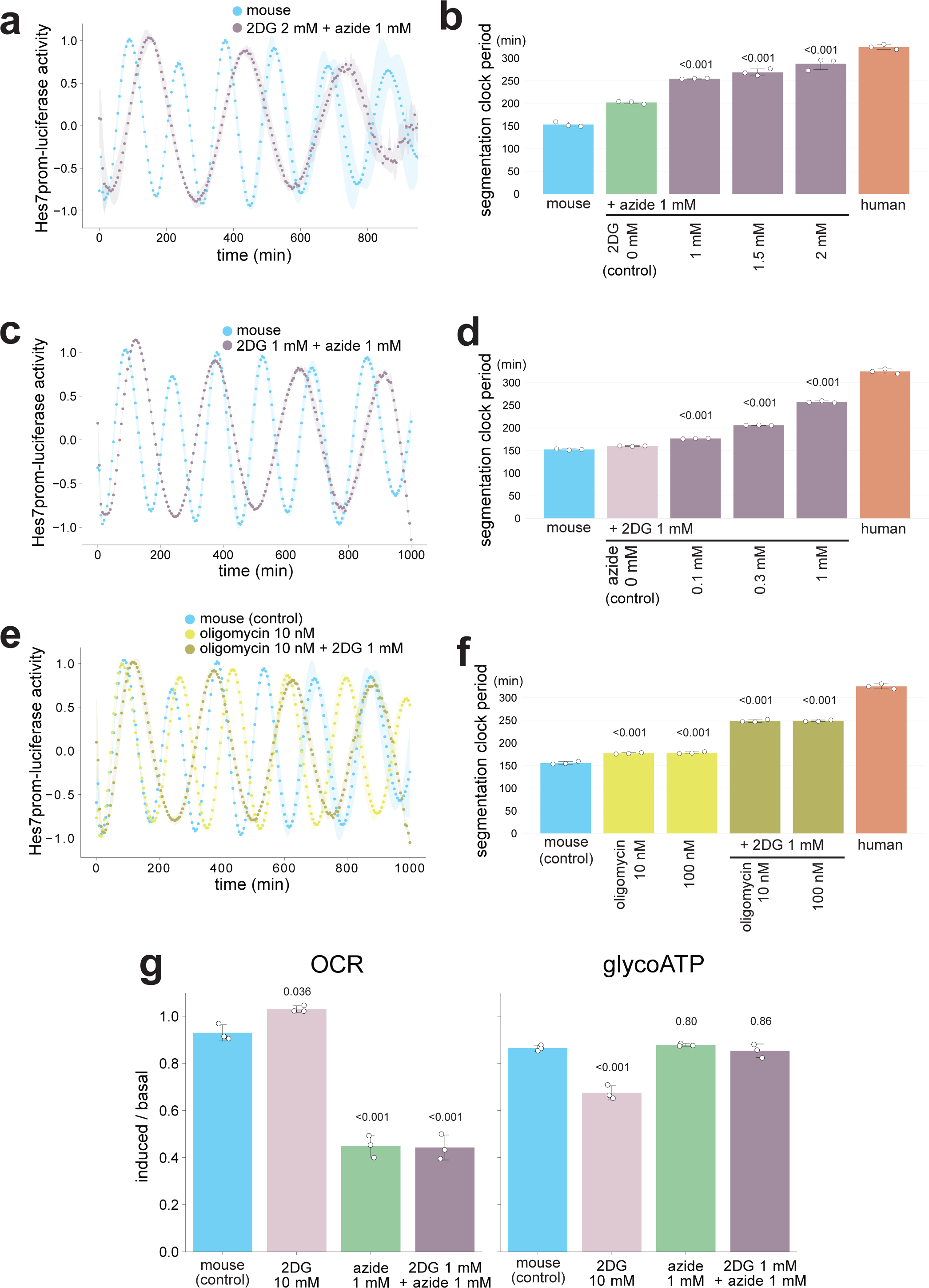
Synergistic effects of metabolic inhibitions on the segmentation clock period. **a,c,e,** Synergistic effects of 2DG, azide, and oligomycin on the segmentation clock period. Mouse PSM cells were pre-treated with the inhibitors for 4 h, and the oscillatory activity of the Hes7 promoter-luciferase reporter was monitored. The signal was detrended and amplitude-normalized. Shading indicates mean ± sd (n = 3). **b,d,f,** Hes7 oscillation periods estimated from a,c,e. Human data are from Fig. 5. **g,** Oxygen consumption rate (OCR) and glycolytic ATP production rate (glycoATP) measured by the Seahorse analyzer. The ratio of induced rates (after adding inhibitors) to basal rates (before adding inhibitors) was calculated from Supplementary Fig. 5. **b,d,f,g,** Graphs indicate mean ± sd (n = 3). P-values are from two-sided Dunnett’s test against the indicated controls.

To assess the effects of the different metabolic inhibitors on the cellular metabolic rates, we measured the oxygen consumption rate (OCR) and glycolytic ATP production rate (glycoATP) (Fig. 3g; Supplementary Fig. 5). The OCR and glycoATP were used as indicators for mitochondrial respiration and glycolytic rates, respectively. Treatment with the ETC inhibitor azide lowered the OCR without influencing the glycoATP (Fig. 3g), as expected. By contrast, the glycolysis inhibitor 2DG lowered the glycoATP and slightly upregulated the OCR, potentially due to a metabolic compensation mechanism (Fig 3g; 10 mM 2DG). Combined treatment with a low concentration of 2DG and azide repressed the OCR to the same extent as azide treatment alone and did not significantly influence the glycoATP (Fig. 3g; 1 mM 2DG + 1 mM azide). In other words, the combined treatment did not exhibit any detectable synergistic effect on the metabolic rates, even though the same combination (1 mM 2DG + 1 mM azide) synergistically extended the segmentation clock period (Fig. 3b,d). These results confirmed the efficacy of the metabolic inhibitors and suggested that the segmentation clock period did not directly correlate with the bio-energetic rates, at least with the OCR or glycoATP.

### The combination of glycolysis and ETC inhibitions simultaneously affects all segmentation clock processes

The notable and synergistic effects of the combined metabolic inhibitions on the segmentation clock period prompted us to investigate their impact on the three key kinetic parameters of Hes7. The combined treatments with 2DG and azide, or 2DG and oligomycin, synergistically decelerated Hes7 protein degradation (Fig. 4a-d; Supplementary Fig. 6). For example, the Hes7 half-life in mouse PSM cells treated with 1 mM 2DG and 1 mM azide (32 ± 1.6 min) was clearly longer than that in untreated cells, while the treatment with the low concentration of 2DG or azide alone did not significantly influence the protein degradation (Fig. 4b). The treatment with 10 nM oligomycin and 1 mM 2DG further extended the Hes7 half-life to 67 ± 4.2 min (Fig. 4d). By contrast, the impact of the combined treatment with 2DG and azide on Hes7 intron delay (19 ± 2.3 min) was similar to that of the azide treatment alone (20 ± 2.0 min in Fig. 2g), and the combination of oligomycin and 2DG did not significantly influence the intron delay (Fig. 4e; Supplementary Fig. 7). The combined treatment with 2DG and azide synergistically extended Hes7 production delay (21 ± 2.4 min; Fig. 4f,g; Supplementary Fig. 8a), while the combination of oligomycin and 2DG showed no significant effect on the production delay (Fig. 4h,i; Supplementary Fig. 8b). These results indicated that combined metabolic inhibitions could yield synergistic effects, especially on Hes7 protein degradation and production delay. It is noteworthy that the combination of glycolysis and ETC inhibitions affected the three key molecular processes simultaneously (1 mM 2DG + 1 mM azide in Fig. 4), fulfilling the criteria as a global modulator for the segmentation clock tempo.

**Figure 4.**
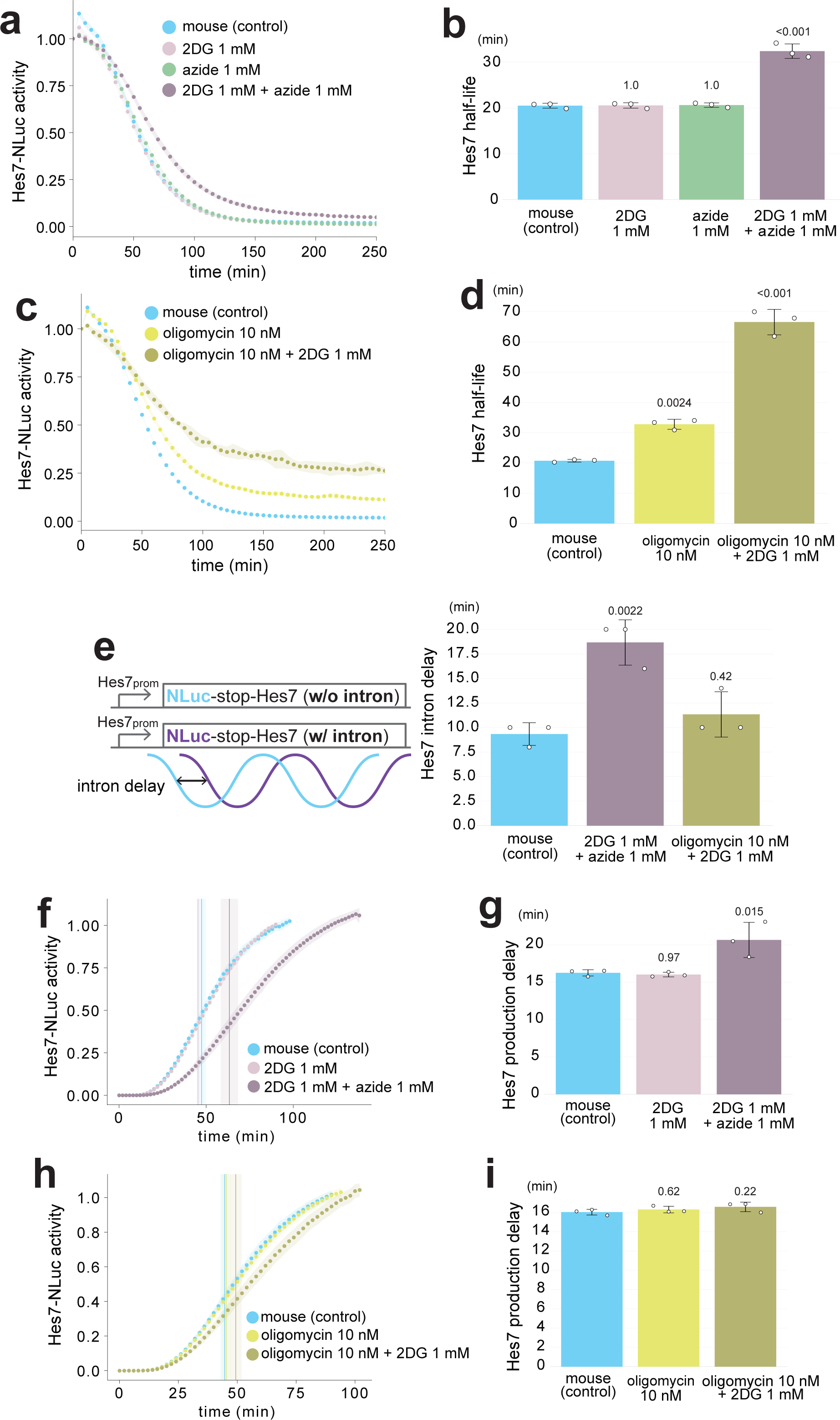
Global effects of combined metabolic inhibitions on segmentation clock processes. **a,c,** Effects of 2DG, azide, and oligomycin on Hes7 protein degradation. Mouse PSM cells were pre-treated with the inhibitors for 4 h before Dox addition, and the decay of the Hes7-NLuc signal was monitored. **b,d,** Hes7 half-lives estimated from a,c and Supplementary Fig. 6. **e,** Effects of 2DG, azide, and oligomycin on Hes7 intron delay. Mouse PSM cells were pre-treated with the inhibitors for 4 h, and the oscillatory activities of the luciferase reporters w/o and w/ Hes7 intron sequences were monitored. The intron delays were estimated from Supplementary Fig. 7. **f,h,** Effects of 2DG, azide, and oligomycin on Hes7 production delay. Mouse PSM cells were pre-treated with the inhibitors for 4 h before Dox addition, and the onset of the Hes7-NLuc signal was monitored. Vertical lines indicate the inflection points. **g,i,** Hes7 production delays estimated from f,h and Supplementary Fig. 8. **a,c,f,h,** Shading indicates mean ± sd (n = 3). **b,d,e,g,i,** Graphs indicate mean ± sd (n = 3). P-values are from two-sided Dunnett’s test against the indicated controls.

### Glycolysis or ETC inhibition selectively affects human segmentation clock processes

We have so far demonstrated the selective effects of distinct metabolic inhibitions on the three key kinetic parameters of the segmentation clock in mouse PSM cells. To shed light on the scaling mechanisms of these parameters across animal species (Fig. 1c), we tested if the human segmentation clock processes show similarly selective responses to metabolic inhibitions. Glycolysis inhibition by 2DG treatment or ETC inhibition by azide treatment extended the human segmentation clock period (Fig. 5a,b). The HES7 protein degradation in human PSM cells was decelerated by 2DG, whereas it was not significantly influenced by azide (Fig. 5c,d; Supplementary Fig. 9). These effects of metabolic inhibitions on the human segmentation clock were consistent with those on the mouse clock. Note, however, that the impacts of 10 mM 2DG on the human segmentation clock appeared slightly milder than those on the mouse clock (compare Fig. 5b,d to Fig. 2c,e). The HES7 intron delay in human PSM cells was notably extended by azide but not by 2DG (Fig. 5e; Supplementary Fig. 10). The HES7 production delay in human PSM cells was extended by 2DG (Fig. 5f,g; Supplementary Fig. 11). Although the production delay was also extended by azide, the extension was negligible. These results obtained in human PSM cells were essentially consistent with the results from mouse PSM cells where 2DG predominantly affected Hes7 protein degradation and production delay whereas azide targeted Hes7 intron delay. As expected, 2DG treatment inhibited glycoATP whereas azide inhibited OCR in human PSM cells (Fig. 5h; Supplementary Fig. 12). Collectively, we concluded that metabolic inhibitions selectively affected the segmentation clock processes in multiple animal species.

**Figure 5.**
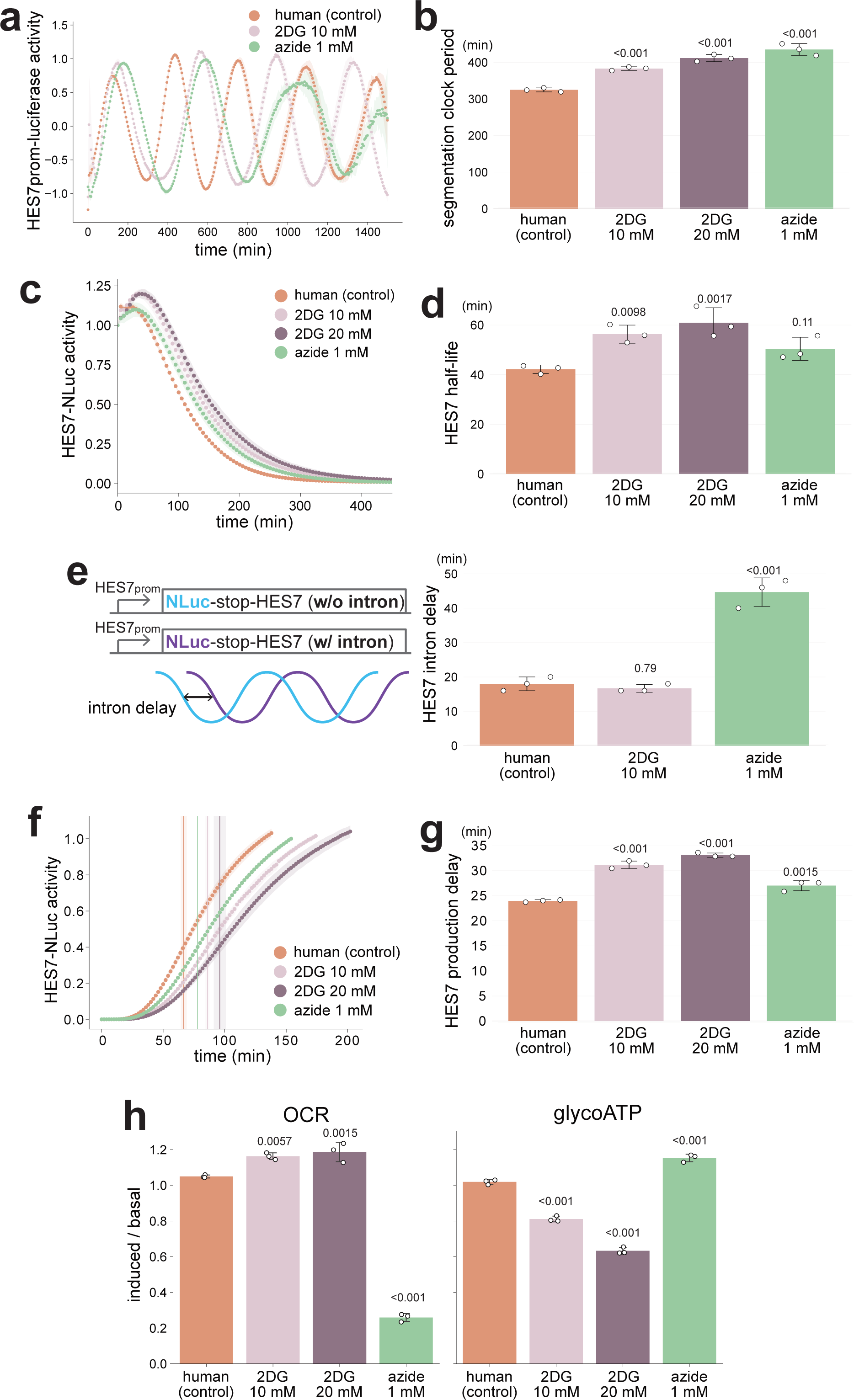
Selective effects of metabolic inhibitions on the human segmentation clock. **a,** Effects of 2DG or azide on the human segmentation clock period. Human PSM cells were treated with the inhibitors from time 0, and the oscillatory activity of the HES7 promoter-luciferase reporter was monitored. The signal was detrended and amplitude-normalized. **b,** HES7 oscillation periods estimated from a. **c,** Effects of 2DG or azide HES7 protein degradation. Human PSM cells were pre-treated with the inhibitors for 5 h before Dox addition, and the decay of the HES7-NLuc signal was monitored. **d,** HES7 half-lives estimated from c and Supplementary Fig. 9. **e,** Effects of 2DG or azide on HES7 intron delay. Human PSM cells were treated with the inhibitors from time 0, and the oscillatory activities of the luciferase reporters w/o and w/ HES7 intron sequences were monitored. The intron delays were estimated from Supplementary Fig. 10. **f,** Effects of 2DG or azide on HES7 production delay. Human PSM cells were pre-treated with the inhibitors for 4 h before Dox addition, and the onset of the HES7-NLuc signal was monitored. Vertical lines indicate the inflection points. **g,** HES7 production delays estimated from f and Supplementary Fig. 11. **h,** Oxygen consumption rate (OCR) and glycolytic ATP production rate (glycoATP) measured by the Seahorse analyzer. The ratio of induced rates (after adding inhibitors) to basal rates (before adding inhibitors) was calculated from Supplementary Fig. 12. **a,c,f,** Shading indicates mean ± sd (n = 3). **b,d,e,g,h,** Graphs indicate mean ± sd (n = 3). P-values are from two-sided Dunnett’s test against the indicated controls.

### Temperature is a global modulator for segmentation clock processes

To contrast the selective modulation by metabolic inhibitions, we employed a temperature shift, a well-established method to modulate the segmentation clock in ectotherms^6,7,37^. In zebrafish, for example, the segmentation clock period changes more than 3 fold across the physiological developmental temperature range^7^: 18.7 min at 30.8°C and 55.4 min at 20.0°C. Consistent with this, we found that decreasing the culture temperature of mouse PSM cells from the standard 37°C to 30°C extended the mouse segmentation clock period (330 ± 3.8 min), mimicking well the human period (Fig. 6a,b). We then examined the effects of the temperature decrease on the three key kinetic parameters of Hes7. Decreasing the temperature decelerated Hes7 protein degradation in mouse PSM cells (Hes7 half-life: 41 ± 2.4 min; Fig. 6c,d; Supplementary Fig. 13a). Moreover, the temperature decrease extended both Hes7 intron delay (35 ± 1.2 min; Fig. 6e; Supplementary Fig. 13b) and production delay (24 ± 1.2 min; Fig. 6f,g; Supplementary Fig. 13c). These results indicated that temperature could act as a global modulator that controls the three key molecular processes simultaneously. However, mice and humans should not have a large difference (such as 7°C) in their body temperatures, and the mouse and human PSM cells cultured in vitro at the same temperature still show differential clock periods. Thus, temperature is a non-physiological global modulator in the context of mammalian segmentation clocks.

**Figure 6.**
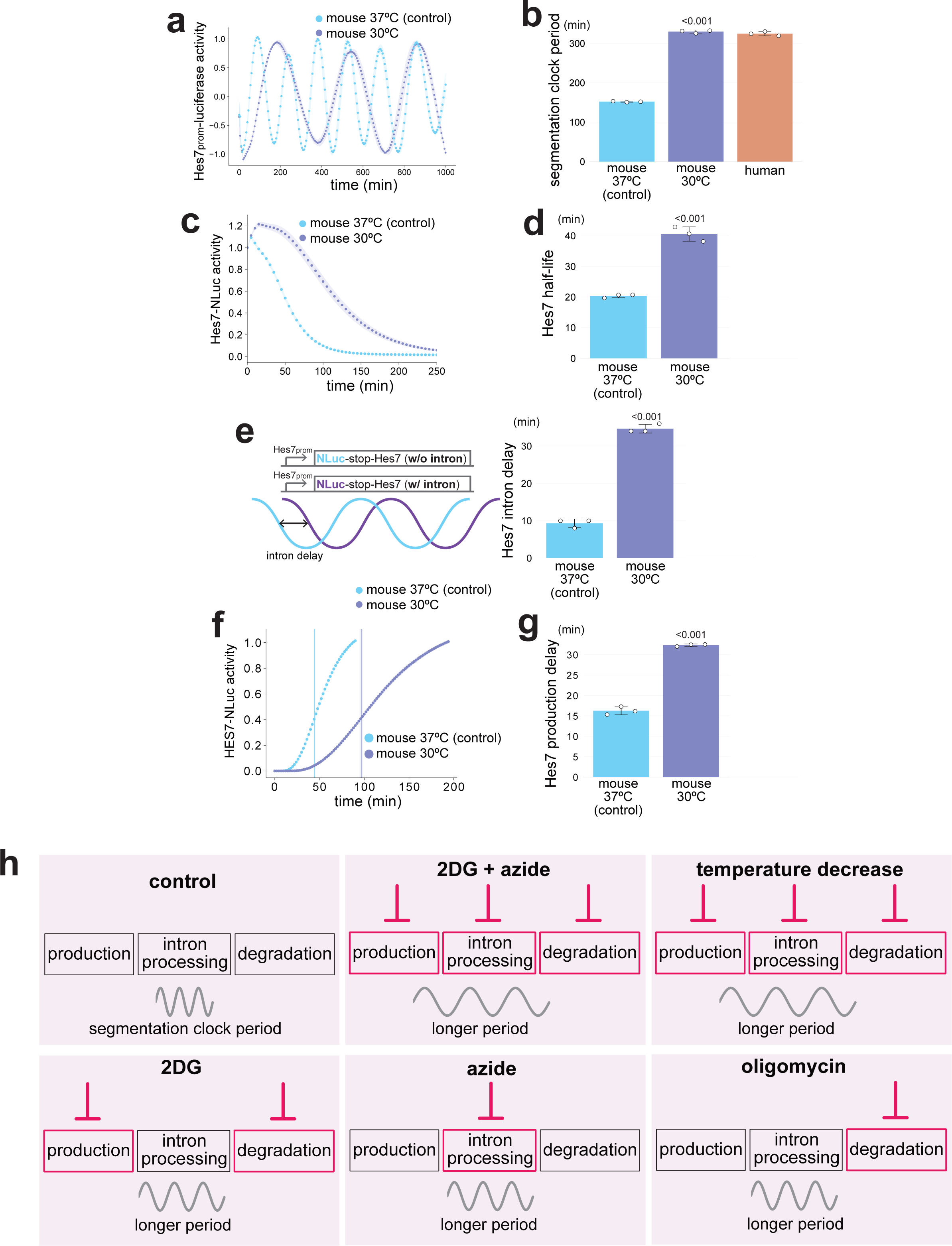
Global effects of temperature change on segmentation clock processes. **a,** Effects of temperature decrease on the segmentation clock period. Mouse PSM cells were pre-incubated at 30°C for 3 h, and the oscillatory activity of the Hes7 promoter-luciferase reporter was monitored. The signal was detrended and amplitude-normalized. **b,** Hes7 oscillation periods estimated from a. **c,** Effects of temperature decrease on Hes7 protein degradation. Mouse PSM cells were pre-incubated at 30°C for 3 h before Dox addition, and the decay of the Hes7-NLuc signal was monitored. **d,** Hes7 half-lives estimated from c and Supplementary Fig. 13a. **e,** Effects of temperature decrease on Hes7 intron delay. Mouse PSM cells were pre-incubated at 30°C for 3 h, and the oscillatory activities of the luciferase reporters w/o and w/ Hes7 intron sequences were monitored. The intron delays were estimated from Supplementary Fig. 13b. **f,** Effects of temperature decrease on Hes7 production delay. Mouse PSM cells were pre-incubated at 30°C for 3 h before Dox addition, and the onset of the Hes7-NLuc signal was monitored. Vertical lines indicate the inflection points. **g,** Hes7 production delays estimated from f and Supplementary Fig. 13c. **h,** Proposed scheme. While both metabolic inhibitions and temperature change extend the segmentation clock period, metabolic inhibitions selectively affect the three key molecular processes: 2DG extends Hes7 production delay and decelerates the protein degradation; azide extends Hes7 intron delay; oligomycin decelerates Hes7 protein degradation. By contrast, the combination of 2DG and azide affects all three processes like temperature change. **a-g,** Mouse 37°C and human data are from Fig. 2 and Fig. 5, respectively. **a,c,f,** Shading indicates mean ± sd (n = 3). **b,d,e,g,** Graphs indicate mean ± sd (n = 3). P-values are from two-sided Student’s t test against the indicated controls.

## Discussion

In this study, we demonstrated the selective effects of metabolic activities on the segmentation clock processes. ETC and ATP synthesis inhibitions exclusively affected Hes7 intron processing and protein degradation processes, respectively, among the three key molecular processes, whereas glycolysis inhibition selectively affected Hes7 protein degradation and production processes (Fig. 6h). Although metabolism had been an attractive candidate for a global modulator of the segmentation clock tempo due to the universal role of energy, distinct metabolic inhibitions were not as universal as temperature change which affected all three processes simultaneously. Instead, the combination of glycolysis and ETC inhibitions affected the three key molecular processes, synergistically extending the segmentation clock period.

How distinct metabolic inhibitions selectively affect the individual molecular processes remains elusive. As bio-energetic rates did not correlate with the segmentation clock period or key kinetic parameters, specific metabolites and metabolic factors that are regulated by distinct metabolic inhibitors, such as glycolytic metabolites and NAD+/NADH^26,36^, may modulate the individual processes. It is also noteworthy that many metabolic enzymes and metabolites have moonlighting functions affecting cellular signaling pathways in addition to the canonical bio-energetic function^34,36,38–40^, and that metabolic inhibitors can perturb other branches of metabolic pathways^41^. Clarifying the effectors of distinct metabolic inhibitions that modulate the three key molecular processes of the segmentation clock serves as an important future research avenue. Importantly, however, even in the presence of multiple moonlighting functions and unexpected side effects in the metabolic pathways, all the metabolic inhibitions employed in this study exhibited selective effects.

As metabolic activities examined in this study are not a global modulator for the three key molecular processes of the segmentation clock, a question remains about how the kinetic parameters of these processes strongly scale with the clock period across multiple animal species (Fig. 1c). One possibility is that the three molecular processes may have been individually fine-tuned by combinations of separate mechanisms, including distinct metabolic activities, throughout evolution. As the segmentation clock operates during phylotypic progression where embryos of related species display the highest degree of morphological and molecular resemblance^42,43^, the three kinetic parameters may be under evolutionary constraints. Different species may employ similar combinations of modulation mechanisms, considering that mouse and human PSM cells exhibited qualitatively similar responses to metabolic inhibitions. By contrast, considering that six animal species show apparently random glycolytic and mitochondrial respiration rates^27^, these species may employ different combinations of modulation mechanisms. Even if there is no singular global modulator for the segmentation clock or developmental tempo, identifying selective modulation mechanisms and their combinations in individual species remains a valuable research avenue. Such investigations will contribute to an evolutionary understanding of developmental tempo and then offer diverse strategies to manipulate it.

## Methods

### Stem cell cultures and PSM cell induction

Mouse EpiSCs (from RIKEN BRC #AES0204)^44^ were maintained on fibronectin-coated dishes with the DMEM-F12 maintenance medium containing 15% Knockout Serum Replacement, Glutamax (2 mM), non-essential amino acids (0.1 mM), β-mercaptoethanol (0.1 mM), Activin A (20 ng/ml), bFGF (10 ng/ml), and IWR-1-endo (2.5 μM). Cells were passaged with ROCK inhibitor Y-27632 (10 μM). The medium was changed every day. For mouse PSM induction, 5 × 10^4^ mouse EpiSCs were seeded on a Matrigel-coated 35 mm dish and cultured in the maintenance medium without IWR-1 for one day. Then the medium was changed into CDMi containing SB431542 (10 μM), CHIR99021 (10 μM), DMH1 (2 μM), and bFGF (20 ng/ml); this medium is hereafter referred to as the SCDF medium. The cells were cultured in the SCDF medium for 30 hours.

Human iPSCs (201B7 line, from RIKEN BRC #HPS0063)^45^ were cultured on Matrigel-coated dishes in the StemFit medium (Ajinomoto). Cells were passaged with ROCK inhibitor Y-27632 (10 μM). Ethical approval for human iPSC usage was granted by Department de Salut de la Generalitat de Catalunya (Carlos III Program). Our human PSM induction protocols have two variations: 1-step and 2-step protocols. For the 1-step protocol, 2 × 10^4^ human iPSCs were seeded on a Matrigel-coated 35 mm dish and cultured for three days. Then the cells were cultured in the SCDF medium for three more days. For the 2-step protocol, 2 × 10^4^ human iPSCs were seeded on a Matrigel-coated 35 mm dish and cultured for three to four days. Then the medium was changed into CDMi containing bFGF (20 ng/ml), CHIR99021 (10 μM), and Activin A (20 ng/ml), and the cells were cultured for one day. The cells were further cultured in the SCDF medium for one more day.

### DNA constructs and reporter cell lines

For the Hes7 oscillation assay, the mHes7 promoter-Firefly luciferase (FLuc)-NLS-PEST-stop-mHes7 (w/o intron) construct was used. For the Hes7 protein degradation assay, the reverse TetOne (rTetOne) promoter-mHes7 (w/o intron)-NLuc construct was used. For the Hes7 intron delay assay, the mHes7 promoter-NLuc-NLS-PEST-stop-mHes7 (w/o intron or w/ intron) construct was introduced into the cell line for the Hes7 oscillation assay. For the Hes7 production assay, the TetOne promoter-mHes7 (w/o intron)-NLuc construct was used. For the measurements in human cells, the constructs derived from hHES7 sequences were used. All components are described previously^25^. All promoters or genes were subcloned into pDONR vector to create entry clones. These entry clones were recombined with a piggyBac vector^46^ (a gift from K. Woltjen) or Tol2 vector^47^ (a gift from K. Kawakami) by using the Multisite Gateway technology (Invitrogen). The constructs were stably introduced into the human iPSCs and mouse EpiSCs by electroporation with a 4D Nucleofector (Lonza) or using lipofectamine (Invitrogen).

### Hes7 oscillation assay

As described previously^27^, the mouse EpiSCs or human iPSCs stably expressing the oscillation reporter were used. After the PSM induction (the 1-step protocol was used for human iPSCs), the medium was changed into the SCDF medium containing a lower dosage of CHIR99021 (1 µM) and D-luciferin to monitor the oscillations of the Hes7 promoter-luciferase reporter signal. The PSM cells were pretreated with metabolic inhibitors 0-4 hours before the medium change, and kept in the medium containing each metabolic inhibitor during the measurement. The luminescence signal of the whole dish was measured at each time point with Kronos Dio Luminometer (Atto). The obtained traces were analyzed with pyBOAT 0.9.12^48^. A threshold of 500 (human and mouse 30°C) or 300 (the rest) min was used for Sinc-detrending and amplitude normalization of the signal. The processed signal was then analyzed using wavelets with periods of 200 - 500 (human and mouse 30°C) or 100 - 300 (the rest) min. A Fourier estimate of the wavelet analysis provided a distribution of periods and their corresponding power. The period with the maximum power for each of the signals was considered.

### Hes7 protein degradation assay

As described previously^27^, the mouse EpiSCs and human iPSCs stably expressing Hes7 degradation reporters were used. PSM cells were induced in the presence of Dox (100 ng/ml). The 2-step protocol was used for human iPSCs. The expression of the NLuc-fusion proteins was initiated by washing out Dox and changing the medium into CDMi containing protected furimazine (Promega). After the NLuc signal was confirmed 4-5 hours later, the expression of the fusion protein was halted by Dox (100 ng/ml) addition, and the decay of NLuc signal was monitored with Kronos Dio luminometer. The metabolic inhibitors were added after washing out Dox. To estimate Hes7 half-life, the slope of log2-transformed data was calculated. A RANSAC algorithm (scikit-learn) was used to find the most linear part of the decay curve.

### Hes7 intron delay assay

As described previously^27^, the mouse EpiSCs and human iPSCs stably expressing Hes7 promoter-NLuc-stop-Hes7 (w/ intron) and Hes7 promoter-FLuc-stop-Hes7 (w/o intron) were used. After PSM cells were induced (the 1-step protocol was used for human iPSCs), the medium was changed into CDMi containing protected furimazine and D-luciferin, and the oscillations of the NLuc and FLuc signals were simultaneously monitored with Kronos Dio luminometer. The PSM cells were pretreated with metabolic inhibitors 0-4 hours before the medium change, and kept in the medium containing each metabolic inhibitor during the measurement. To estimate Hes7 intron delay, the oscillation phase difference between the reporter w/o intron and the one w/ intron was estimated by calculating their cross-correlation with Python (SciPy). To normalize the difference in the maturation/degradation time between NLuc and FLuc, cells containing the Hes7 promoter-NLuc-stop-Hes7 (w/o intron) and Hes7 promoter-FLuc-stop-Hes7 (w/o intron) constructs were also used, and the phase difference between the NLuc and FLuc reporters was subtracted from that between the w/o intron and w/ intron reporters.

### Hes7 production delay assay

As described previously^25^, the mouse EpiSCs and human iPSCs stably expressing the production delay assay reporters were used. After PSM cells were induced in the absence of Dox, the medium was changed into CDMi containing protected furimazine and the metabolic inhibitors. Four hours after the medium change, the expression of the fusion protein was initiated by Dox (100 ng/ml), and the onset of NLuc signal was monitored with Kronos Dio luminometer. To estimate Hes7 production delay, the following model was considered.

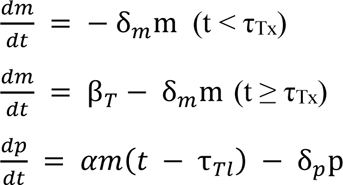

where m and p are the concentrations of Hes7 mRNA and protein, δ_m_ and δ_p_ are the degradation rates of mRNA and protein, β_*T*_ is the transcription rate of the TetOne promoter, α is the translation rate, and τ_Tx_ and τ_Tl_ are the transcription and translation delays.

The solution of this is

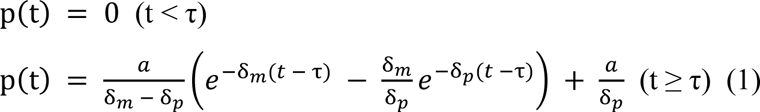

where *τ* = τ_*Tx*_ + τ_*Tl*_, and *a* = *α*β_*T*_/δ_*m*_.

δ_p_ was estimated in the degradation assay. The production delay τ, together with δ_m_ and a, was estimated by fitting the data from the production delay assay to equation (1) with Python (SciPy)’s basin-hopping algorithm. Data points within 2×(duration to reach the inflection point) were used for fitting, and the inflection point was determined by calculating the 2nd derivatives of the data.

### Seahorse metabolic rate measurement

As described previously^27^, PSM cells were re-seeded on a fibronectin-coated Seahorse plate (Agilent) at a density of 9.1 × 10^5^ cells/cm^2^ (mouse assay) or 7.3 × 10^5^ cells/cm^2^ (human assay) in 100 μl of Seahorse XF DMEM (Agilent) supplemented with glucose (20 mM), pyruvate (1 mM), and glutamine (2 mM). Cells were allowed to attach at RT for 10 min and then transferred to a 37°C incubator without CO_2_ for 10 min. After that time, 400 μl of Seahorse XF DMEM medium at 37°C was added carefully to each well without disturbing the attached cells for a total of 500 μl. Cells were incubated at 37°C without CO2 for 30 more min. For the real-time ATP rate assay (Agilent), oligomycin (1 μM), rotenone (0.5 μM), and antimycin A (0.5 μM) were used. MitoATP could not be calculated as a parameter to compare across conditions since azide was added at the beginning in some conditions. The Wave Desktop and online app provided by the manufacturer were used for analysis.

## Acknowledgments

We are grateful to Kristina Stapornwongkul, Hidenobu Miyazawa, and Shuting Xu for their comments on the manuscript. This work was supported by EMBL; the Deutsche Forschungsgemeinschaft (DFG, German Research Foundation) under Germany’s Excellence Strategy - EXC 2068 - 390729961 - Cluster of Excellence Physics of Life of TU Dresden; the European Research Council (ERC) under the European Union’s Horizon 2020 research and innovation program (grant agreement No. 101002564) (to M.E.); PRESTO (grant number JP20332265) from Japan Science and Technology Agency (JST) (to M.M.); the Boehringer Ingelheim Fonds (BIF) PhD fellowship (to J.L.); M.E. is supported by the Alexander von Humboldt Foundation in the framework of the Alexander von Humboldt Professorship endowed by the Federal Ministry of Education and Research. The human iPSC and mouse EpiSC lines were provided by the RIKEN BRC through the National BioResource Project of the MEXT, Japan.

## Author contributions

M.M. and M.E. designed the work and wrote the manuscript. M.M. performed the experiments and analyzed the data. J.L. analyzed the data. All authors contributed to the manuscript and approved the final version.

---

**Supplementary Figure 1 Protein degradation assay a,**

Schematic representation of the rTetOne promoter-Hes7-NLuc reporter. Upon Dox addition at time 0, rTetOne promoter is repressed, and the transcription of Hes7-NLuc is halted. By monitoring the decay of the Hes7-NLuc signal, the protein degradation rate is estimated. **b**, Raw data and fitting of the Hes7 protein degradation assay shown in Fig. 2e. Dashed lines indicate the most linear region considered by the RANSAC algorithm for the fitting. Slope of the fitted line is shown, and it was converted to the half-life using the equation: half-life = −1/slope.

**Supplementary Figure 2 Intron delay assay a,**

Schematic representation of the intron delay assay. As explained in Fig. 2f, Hes7 intron delay is defined as the oscillation phase difference between two reporters without (w/o) and with (w/) Hes7 intron sequences, namely the phase difference between NLuc-Hes7 (w/o intron) (blue) and NLuc-Hes7 (w/ intron) (purple) reporters. However, since different spectra of NLuc and FLuc are required to simultaneously monitor the oscillatory activities of two reporters, the FLuc-Hes7 (w/o intron) reporter (orange) is also required. Step 1 (left panel): To normalize the maturation/degradation time difference between NLuc and FLuc proteins, the phase difference between blue and orange lines is calculated. Step 2 (right panel): The phase difference between purple and orange lines is calculated. Step 3: The intron delay (the phase difference between blue and purple lines) is calculated by subtracting the phase difference in Step 1 from that in Step 2. **b,** Raw data of the Hes7 intron delay assay shown in Fig. 2g. Shading indicates mean ± sd (n = 3). Top: Oscillatory activities of two reporters were monitored simultaneously. The signal was detrended and amplitude-normalized. Bottom: Cross-correlation of the two reporters. The peak of the cross-correlation was used to calculate the oscillation phase difference of the two reporters. The intron delay (the phase difference between blue and purple lines) was estimated by subtracting the phase difference between blue and orange lines from that between purple and orange lines.

**Supplementary Figure 3 Production delay assay a,**

Schematic representation of the TetOne promoter-Hes7 reporter. Upon Dox addition at time 0, TetOne promoter is activated, and the transcription of Hes7-NLuc is initiated. By monitoring the onset of the Hes7-NLuc signal, the production delay is estimated. **b,** Raw data and fitting of the Hes7 production delay assay shown in Fig. 2i. The data within 2×(duration to reach the inflection point) were used for fitting, and the inflection point was determined by calculating the 2nd derivatives of the data.

**Supplementary Figure 4 Effects of metabolic inhibitions on raw oscillatory signals**

Panel a, b, and c represent the raw data of the oscillatory signals shown in Fig. 2b, 3a, and 3e, respectively. Shading indicates mean ± sd (n = 3). The first peak of the oscillatory signal of the control sample was set to 1.

**Supplementary Figure 5 Metabolic rate measurement**

Raw data of the oxygen consumption rate (OCR) and proton efflux rate (PER) measured over the course of the Seahorse real-time ATP rate assay shown in Fig. 3g. Metabolic inhibitors (2DG and azide), oligomycin, and rotenone + antimycin A (rot + AA) were added at the marked time points. Error bars indicate mean ± sd.

**Supplementary Figure 6 Protein degradation assay with combined metabolic inhibitions**

Raw data and fitting of the Hes7 protein degradation assay shown in Fig. 4b,d. Dashed lines indicate the most linear region considered by the RANSAC algorithm for the fitting. Slope of the fitted line is shown, and it was converted to the half-life using the equation: half-life = −1/slope.

**Supplementary Figure 7 Intron delay assay with combined metabolic inhibitions**

Raw data of the Hes7 intron delay assay shown in Fig. 4e. Shading indicates mean ± sd (n = 3). Top: Oscillatory activities of two reporters were monitored simultaneously. The signal was detrended and amplitude-normalized. Bottom: Cross-correlation of the two reporters. The peak of the cross-correlation was used to calculate the oscillation phase difference of the two reporters. The intron delay (the phase difference between blue and purple lines) was estimated by subtracting the phase difference between blue and orange lines from that between purple and orange lines.

**Supplementary Figure 8 Production delay assay with combined metabolic inhibitions**

Raw data and fitting of the Hes7 production delay assay shown in Fig. 4g,i. The data within 2×(duration to reach the inflection point) were used for fitting, and the inflection point was determined by calculating the 2nd derivatives of the data.

**Supplementary Figure 9 Protein degradation assay in human PSM cells**

Raw data and fitting of the HES7 protein degradation assay shown in Fig. 5d. Dashed lines indicate the most linear region considered by the RANSAC algorithm for the fitting. Slope of the fitted line is shown, and it was converted to the half-life using the equation: half-life = −1/slope.

**Supplementary Figure 10 Intron delay assay in human PSM cells**

Raw data of the HES7 intron delay assay shown in Fig. 5e. Shading indicates mean ± sd (n = 3). Top: Oscillatory activities of two reporters were monitored simultaneously. The signal was detrended and amplitude-normalized. Bottom: Cross-correlation of the two reporters. The peak of the cross-correlation was used to calculate the oscillation phase difference of the two reporters. The intron delay (the phase difference between blue and purple lines) was estimated by subtracting the phase difference between blue and orange lines from that between purple and orange lines.

**Supplementary Figure 11 Production delay assay in human PSM cells**

Raw data and fitting of the HES7 production delay assay shown in Fig. 5g. The data within 2×(duration to reach the inflection point) were used for fitting, and the inflection point was determined by calculating the 2nd derivatives of the data.

**Supplementary Figure 12 Metabolic rate measurement in human PSM cells**

Raw data of the oxygen consumption rate (OCR) and proton efflux rate (PER) measured over the course of the Seahorse real-time ATP rate assay shown in Fig. 5h. Metabolic inhibitors (2DG and azide), oligomycin, and rotenone + antimycin A (rot + AA) were added at the marked time points. Error bars indicate mean ± sd.

**Supplementary Figure 13 Temperature change**

Mouse PSM cells were incubated at 30°C instead of the standard 37°C for each assay. **a,** Raw data and fitting of the Hes7 protein degradation assay shown in Fig. 6d. Dashed lines indicate the most linear region considered by the RANSAC algorithm for the fitting. Slope of the fitted line is shown, and it was converted to the half-life using the equation: half-life = −1/slope. **b,** Raw data of the Hes7 intron delay assay shown in Fig. 6e. Shading indicates mean ± sd (n = 3). Top: Oscillatory activities of two reporters were monitored simultaneously. The signal was detrended and amplitude-normalized. Bottom: Cross-correlation of the two reporters. The peak of the cross-correlation was used to calculate the oscillation phase difference of the two reporters. The intron delay (the phase difference between blue and purple lines) was estimated by subtracting the phase difference between blue and orange lines from that between purple and orange lines. **c,** Raw data and fitting of the Hes7 production delay assay shown in Fig. 6g. The data within 2×(duration to reach the inflection point) were used for fitting, and the inflection point was determined by calculating the 2nd derivatives of the data.

