## Supplementary Figures for "Metabolic activities are selective modulators for individual segmentation clock processes"

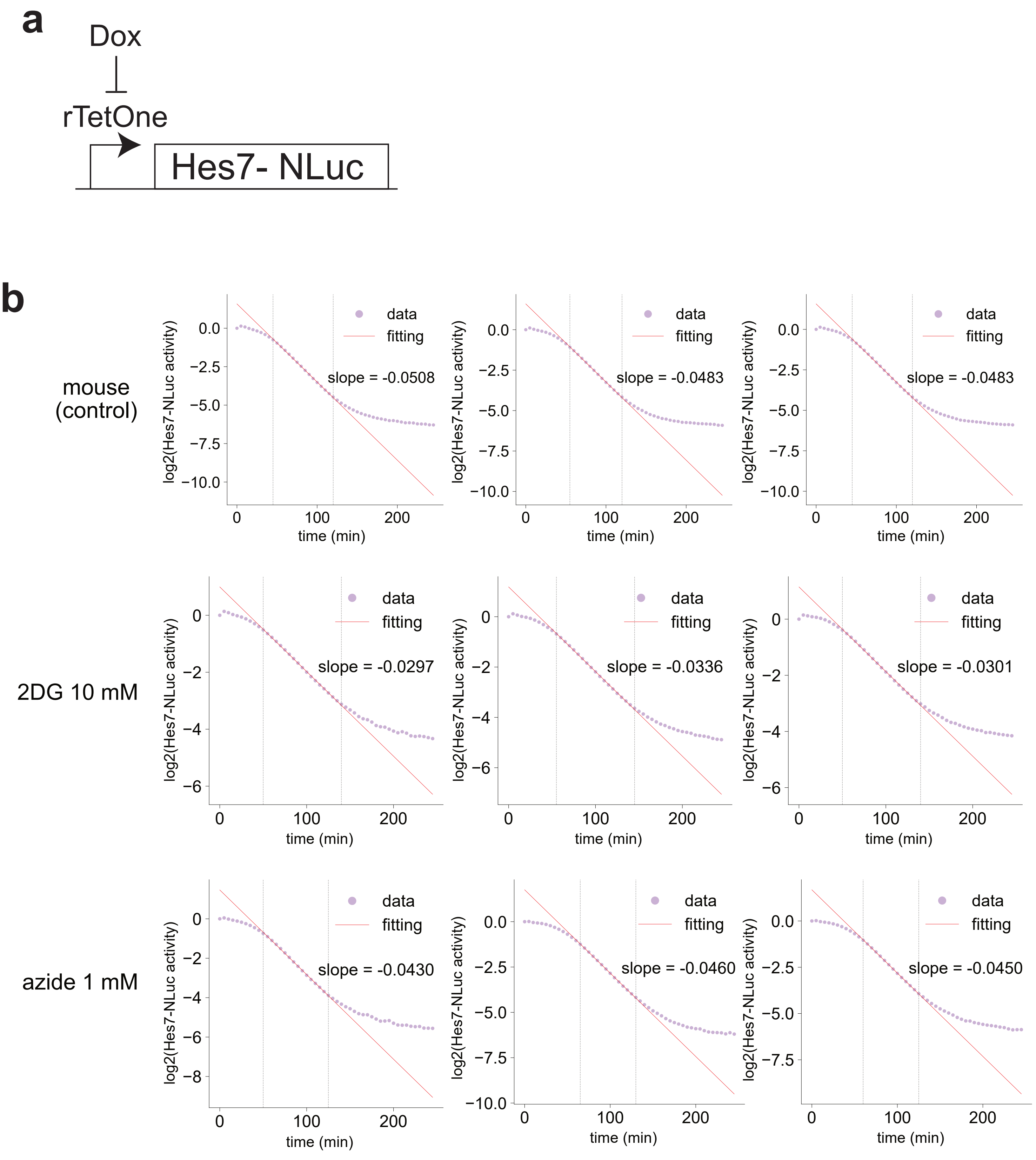
a
Dox
rTetOne
Hes7- NLuc
b
mouse (control)
2DG 10 mM
azide 1 mM
data
fitting
slope = -0.0508
slope = -0.0483
slope = -0.0483
slope = -0.0297
slope = -0.0336
slope = -0.0301
slope = -0.0430
slope = -0.0460
slope = -0.0450
log2(Hes7-NLuc activity)
time (min)

Sup. Fig. 1

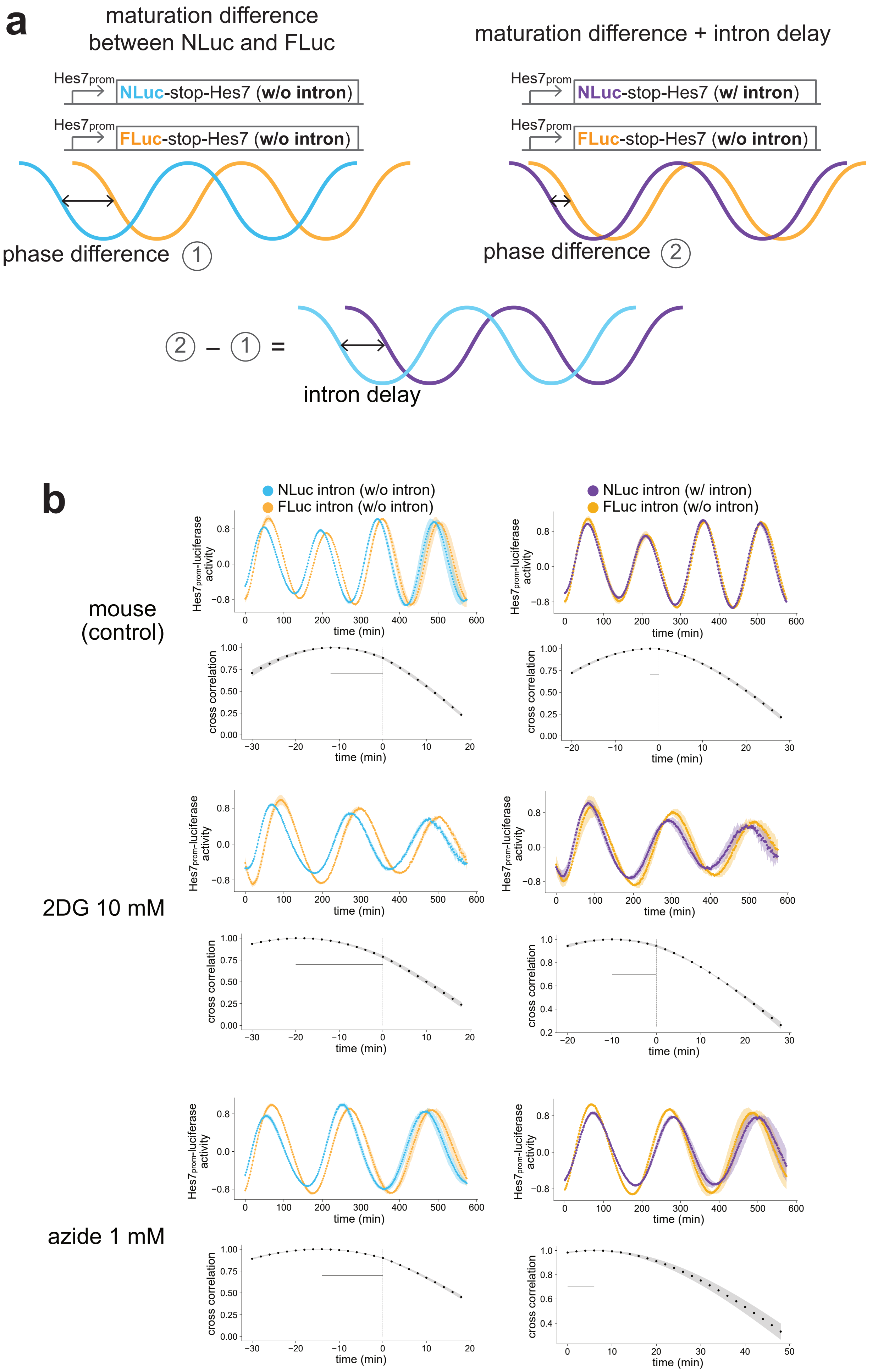
a
maturation difference between NLuc and FLuc
Hes7prom
NLuc-stop-Hes7 (w/o intron)
FLuc-stop-Hes7 (w/o intron)
phase difference 1
maturation difference + intron delay
NLuc-stop-Hes7 (w/ intron)
phase difference 2
2 − 1 =
intron delay
b
NLuc intron (w/o intron)
FLuc intron (w/o intron)
NLuc intron (w/ intron)
mouse (control)
2DG 10 mM
azide 1 mM
Hes7prom-luciferase activity
time (min)
cross correlation

Sup. Fig. 2

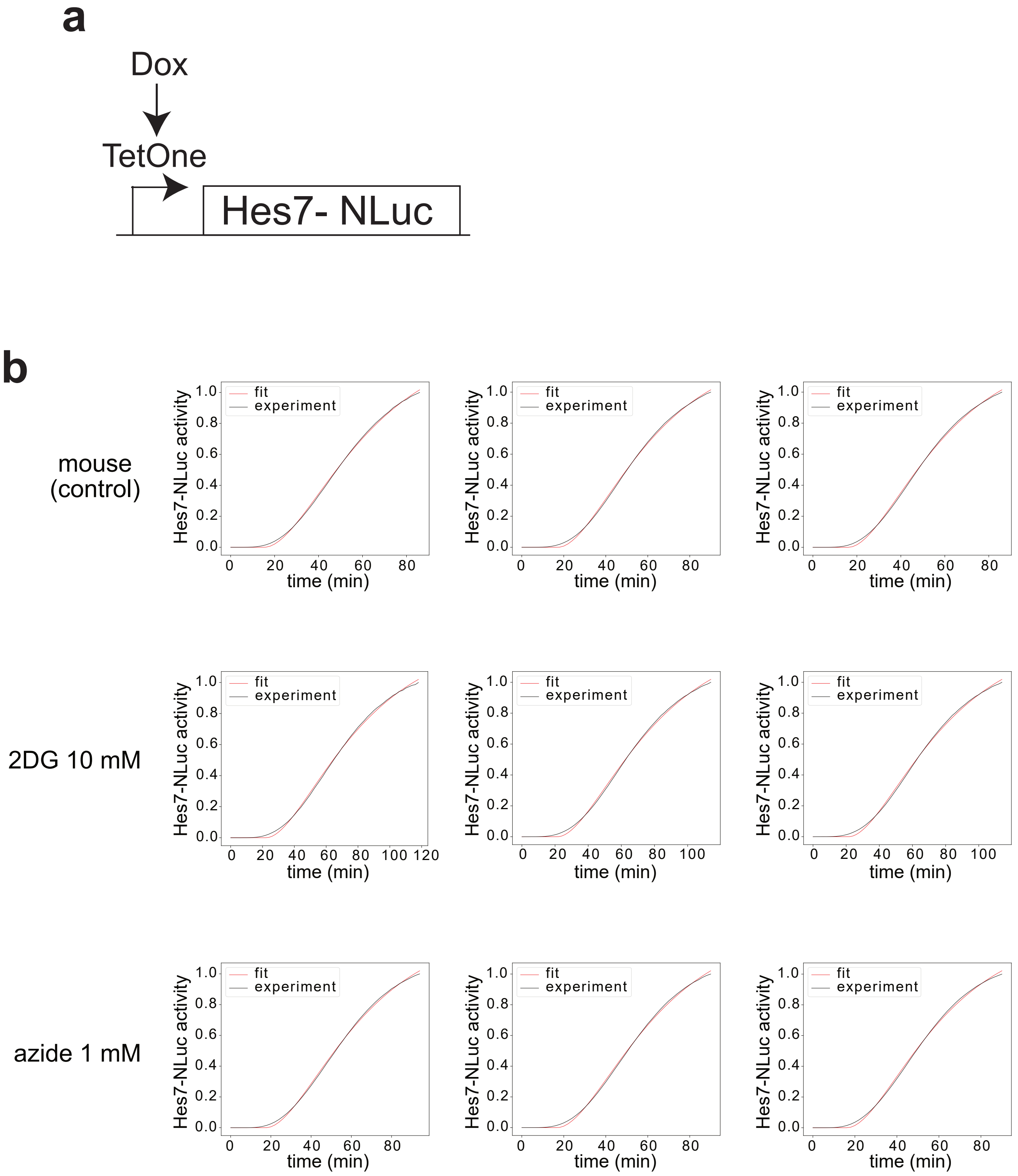
a
Dox
TetOne
Hes7- NLuc
b
mouse (control)
2DG 10 mM
azide 1 mM
fit
experiment
Hes7-NLuc activity
time (min)

Sup. Fig. 3

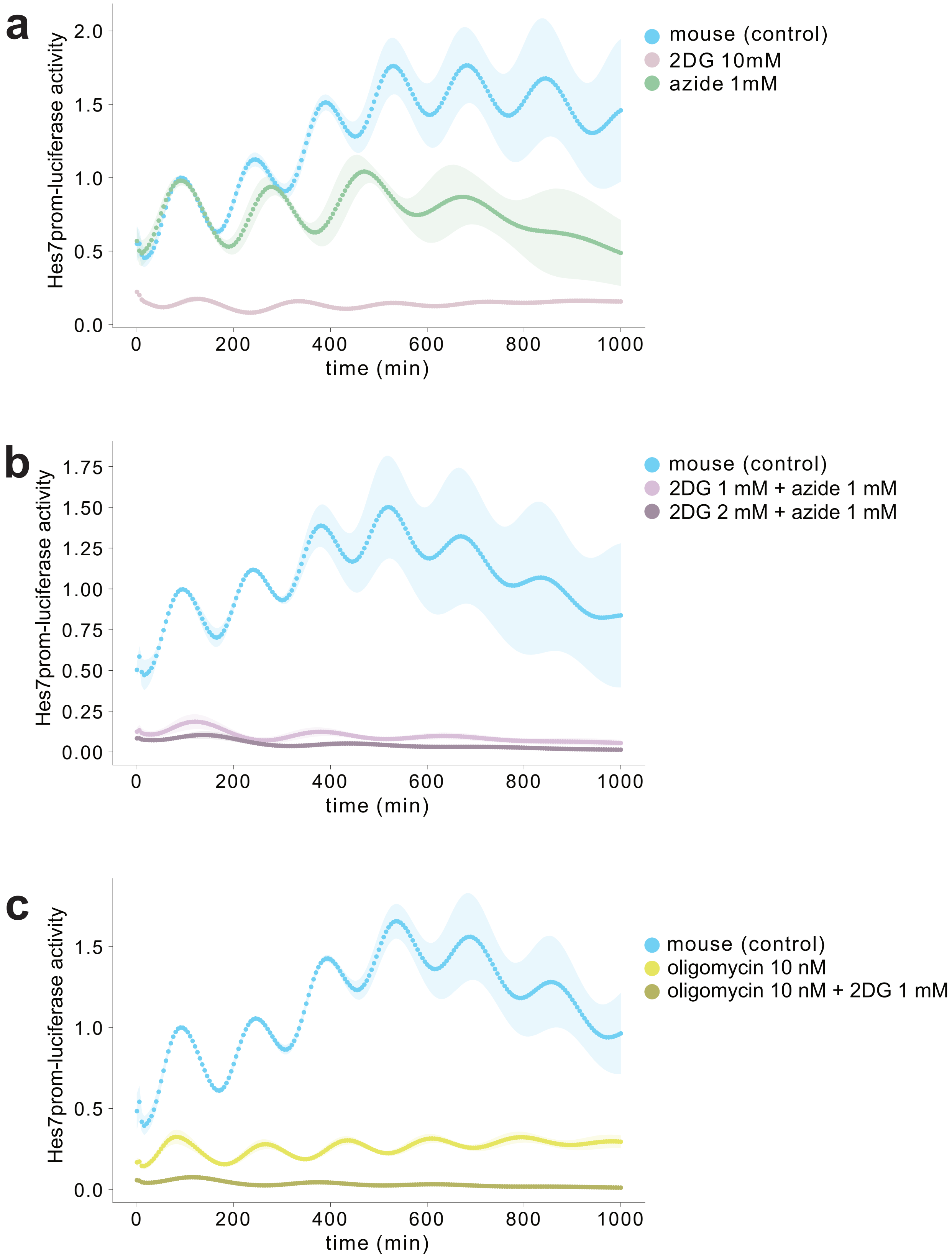
a
b
c
Hes7prom-luciferase activity
time (min)
0
200
400
600
800
1000
0.0
0.5
1.0
1.5
2.0
0.00
0.25
0.50
0.75
1.00
1.25
1.50
1.75
mouse (control)
2DG 10mM
azide 1mM
2DG 1 mM + azide 1 mM
2DG 2 mM + azide 1 mM
oligomycin 10 nM
oligomycin 10 nM + 2DG 1 mM

Sup. Fig. 4

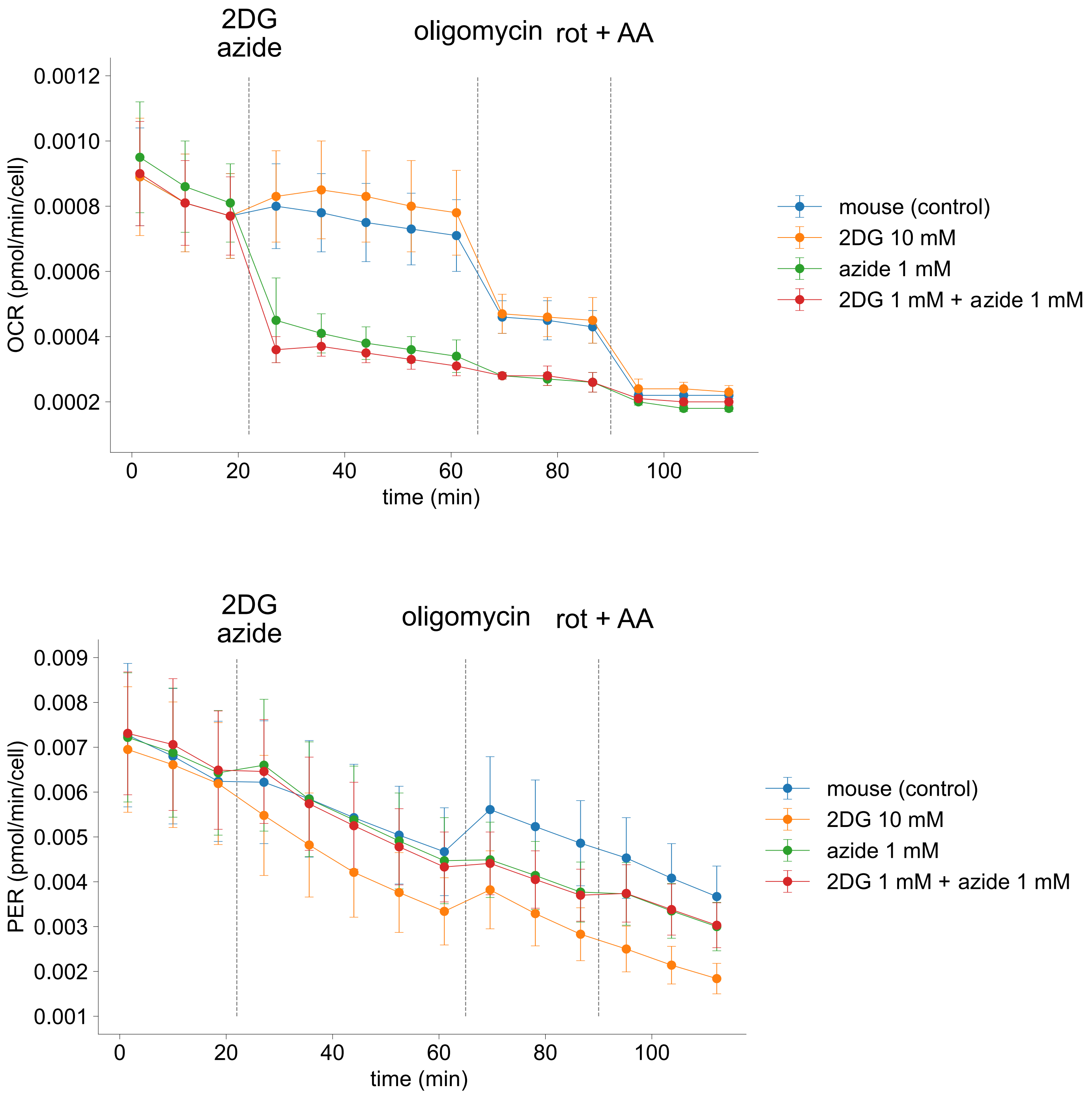
2DG
azide
oligomycin
rot + AA
OCR (pmol/min/cell)
0.0012
0.0010
0.0008
0.0006
0.0004
0.0002
0
20
40
60
80
100
time (min)
mouse (control)
2DG 10 mM
azide 1 mM
2DG 1 mM + azide 1 mM
2DG
azide
oligomycin
rot + AA
PER (pmol/min/cell)
0.009
0.008
0.007
0.006
0.005
0.004
0.003
0.002
0.001
0
20
40
60
80
100
time (min)
mouse (control)
2DG 10 mM
azide 1 mM
2DG 1 mM + azide 1 mM

Sup. Fig. 5

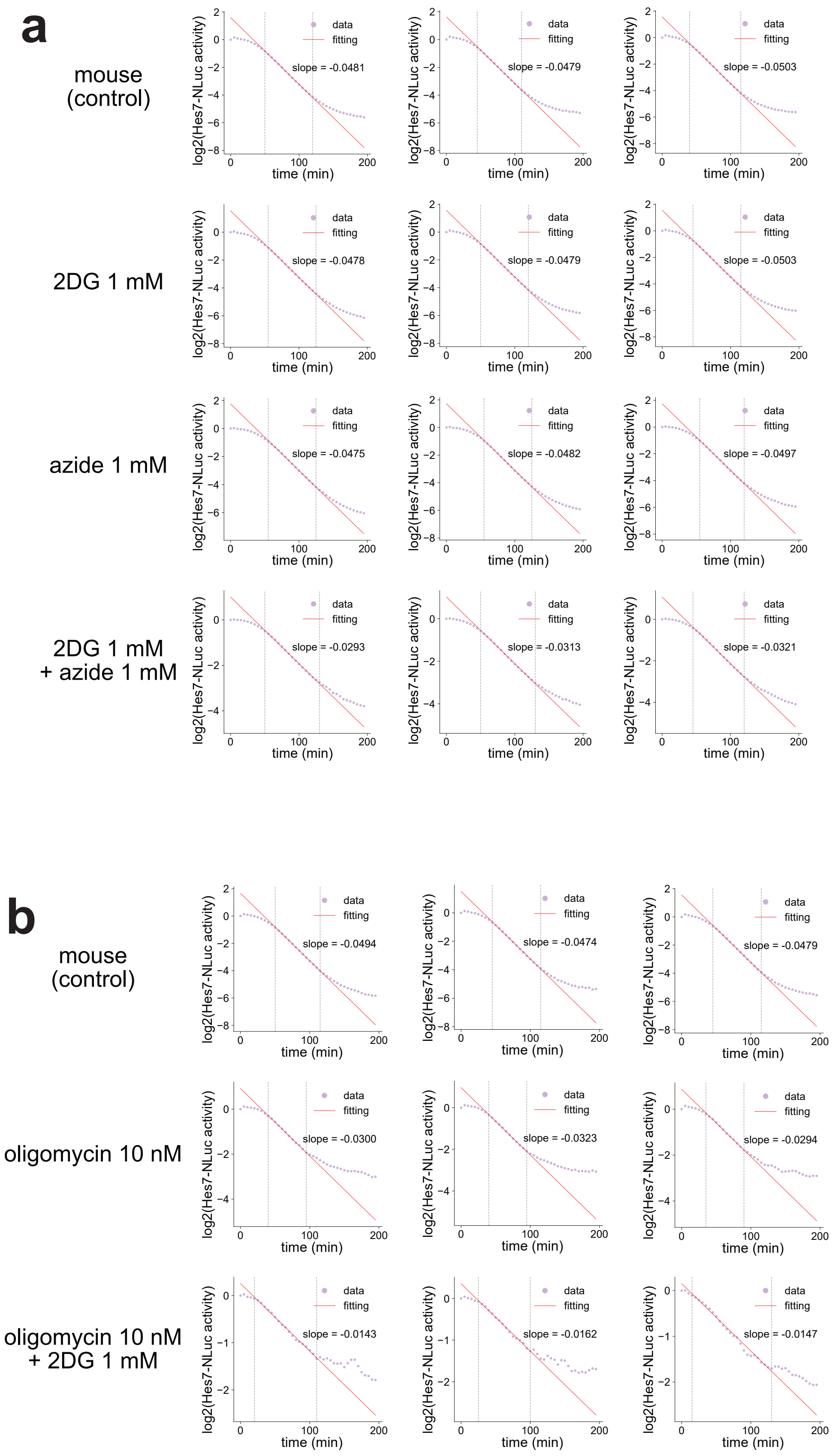
a
mouse
(control)
2DG 1 mM
azide 1 mM
2DG 1 mM
+ azide 1 mM
log2(Hes7-NLuc activity)
time (min)
data
fitting
slope = -0.0481
slope = -0.0479
slope = -0.0503
slope = -0.0478
slope = -0.0479
slope = -0.0503
slope = -0.0475
slope = -0.0482
slope = -0.0497
slope = -0.0293
slope = -0.0313
slope = -0.0321
b
mouse
(control)
oligomycin 10 nM
oligomycin 10 nM
+ 2DG 1 mM
slope = -0.0494
slope = -0.0474
slope = -0.0479
slope = -0.0300
slope = -0.0323
slope = -0.0294
slope = -0.0143
slope = -0.0162
slope = -0.0147

Sup. Fig. 6

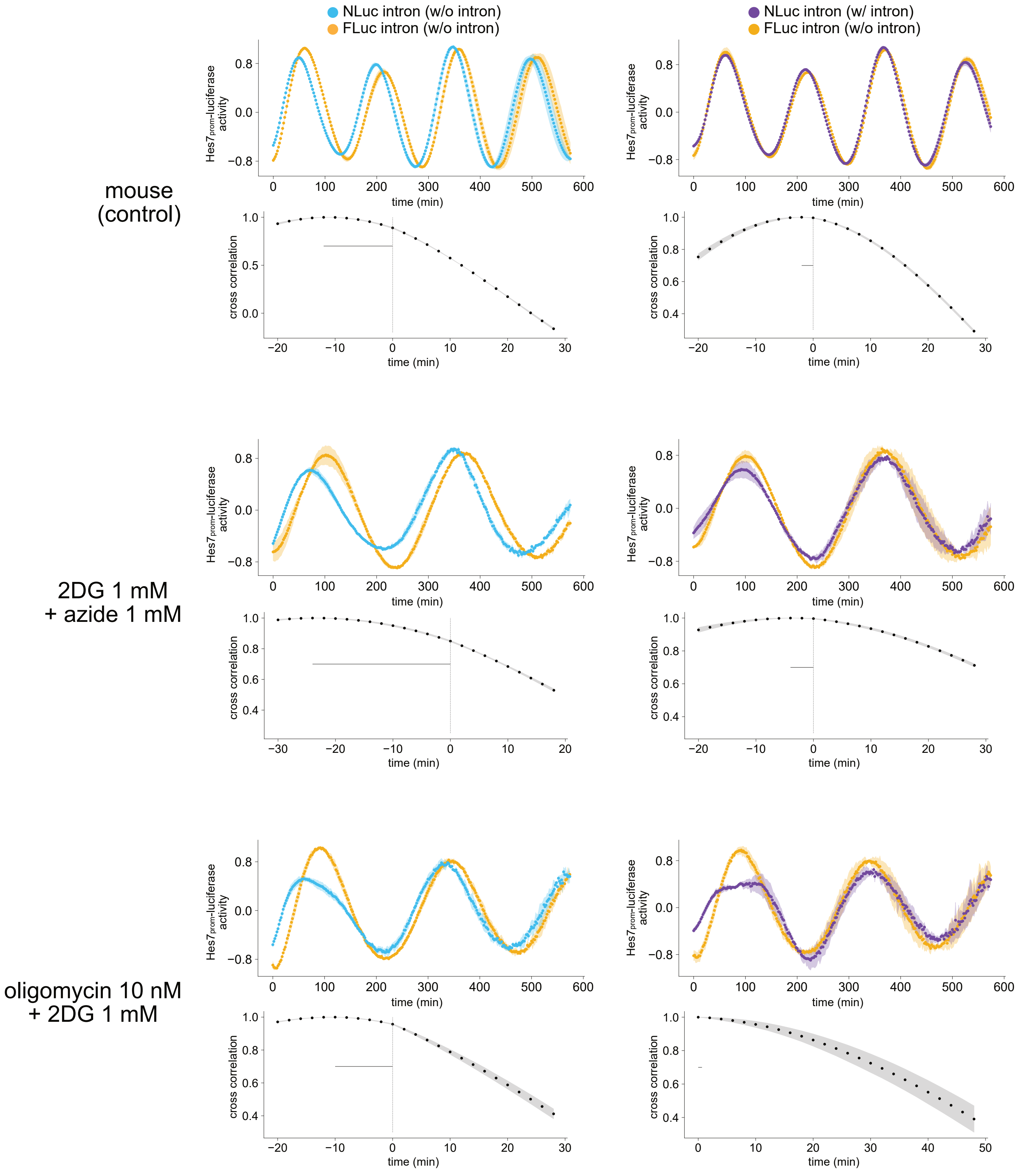
NLuc intron (w/o intron)
FLuc intron (w/o intron)
NLuc intron (w/ intron)
FLuc intron (w/o intron)
mouse (control)
2DG 1 mM + azide 1 mM
oligomycin 10 nM + 2DG 1 mM
Hes7prom-luciferase activity
cross correlation
time (min)

Sup. Fig. 7

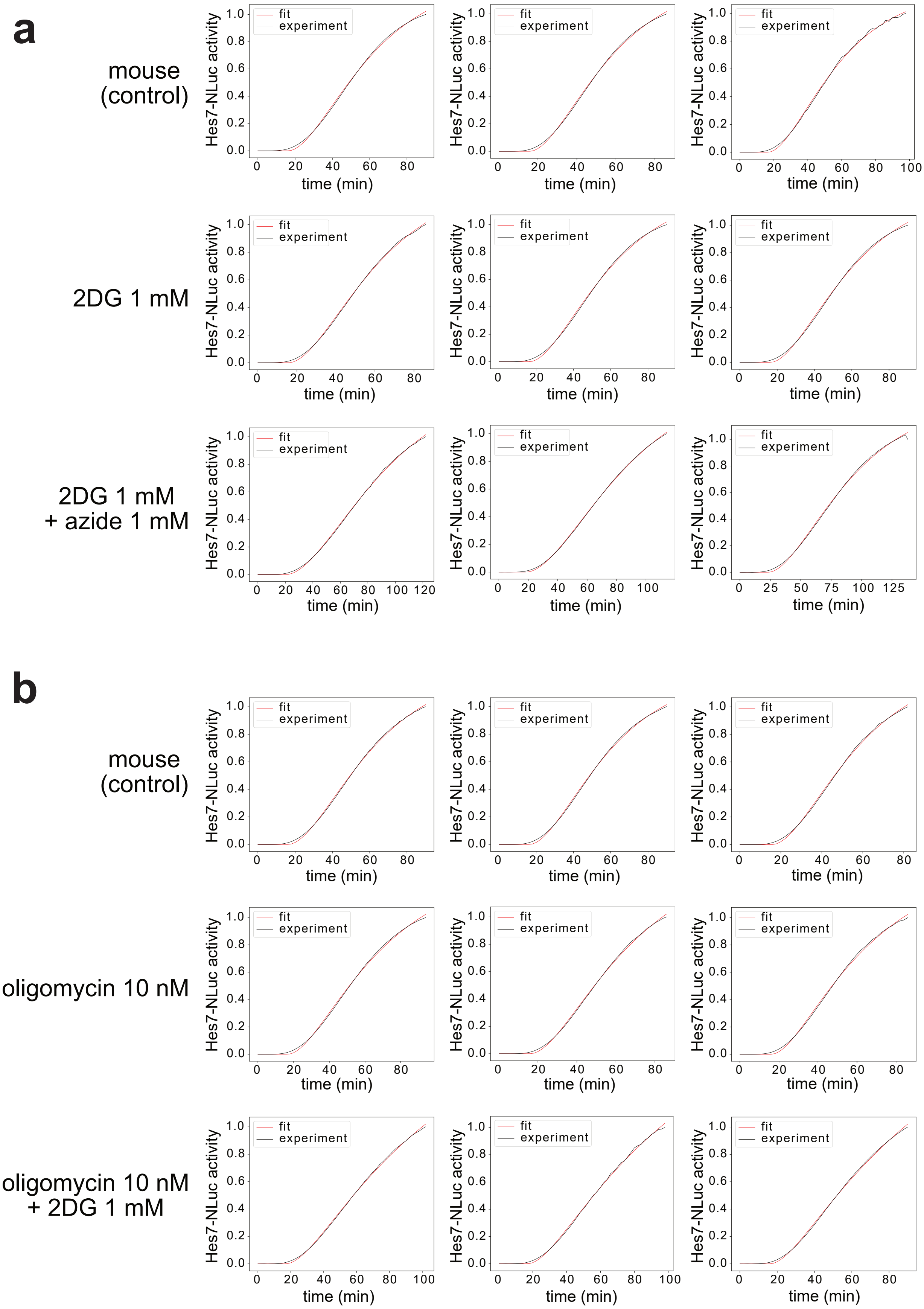
a
mouse (control)
2DG 1 mM
2DG 1 mM + azide 1 mM
b
mouse (control)
oligomycin 10 nM
oligomycin 10 nM + 2DG 1 mM
Hes7-NLuc activity
time (min)
fit
experiment

Sup. Fig. 8

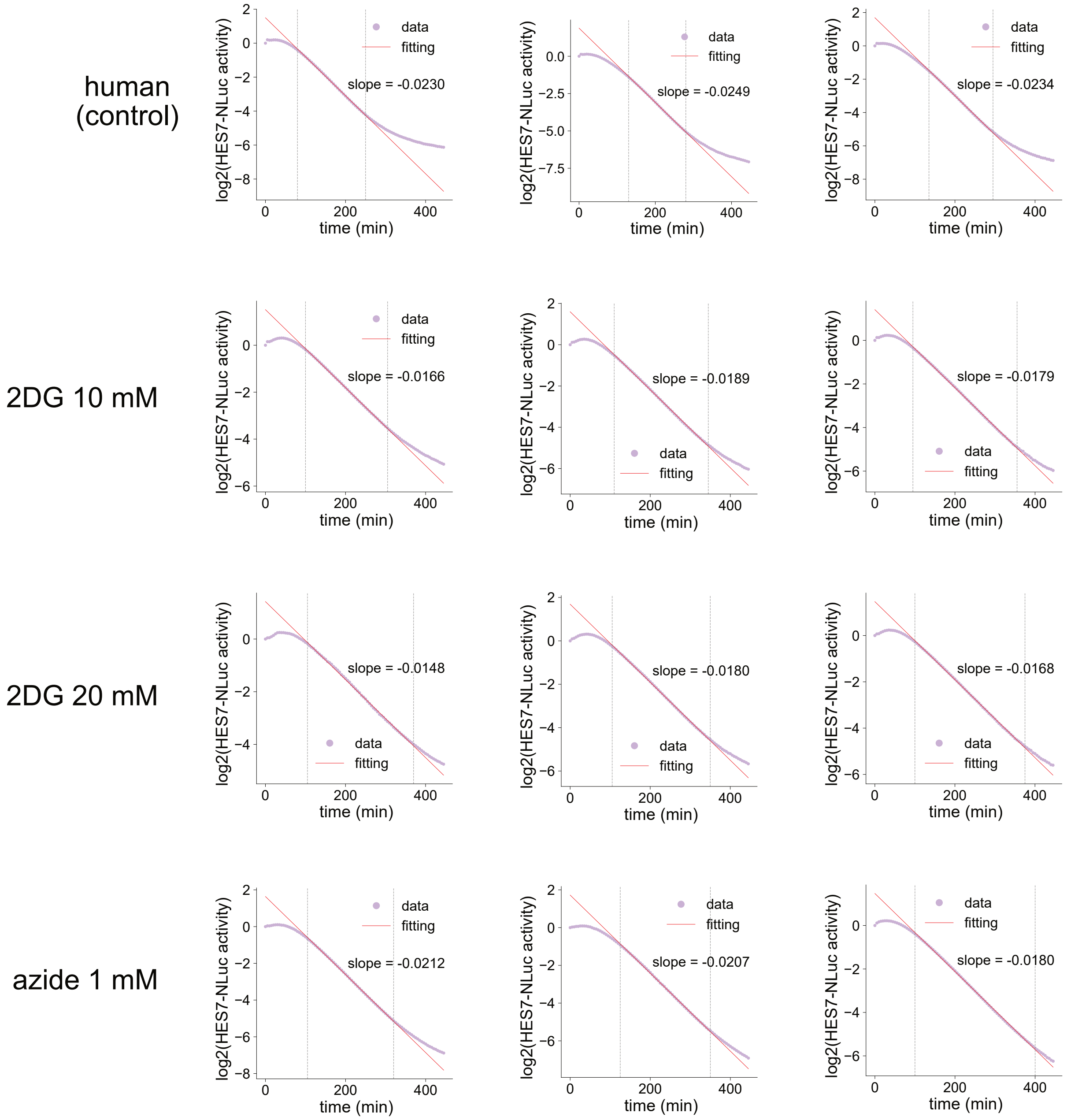
human
(control)
2DG 10 mM
2DG 20 mM
azide 1 mM
log2(HES7-NLuc activity)
time (min)
data
fitting
slope = -0.0230
slope = -0.0249
slope = -0.0234
slope = -0.0166
slope = -0.0189
slope = -0.0179
slope = -0.0148
slope = -0.0180
slope = -0.0168
slope = -0.0212
slope = -0.0207
slope = -0.0180

Sup. Fig. 9

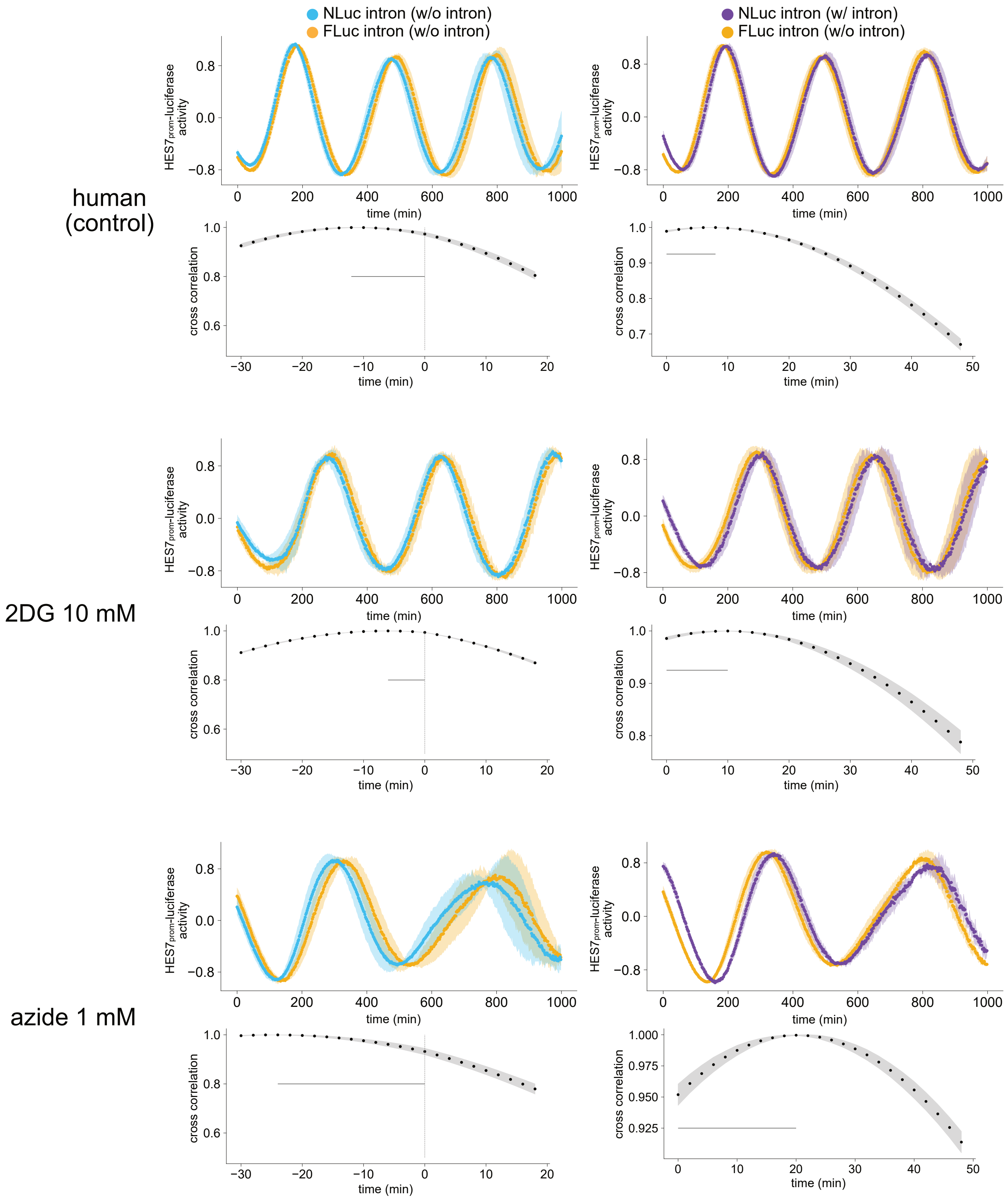

NLuc intron (w/o intron)
FLuc intron (w/o intron)
NLuc intron (w/ intron)
FLuc intron (w/o intron)
human
(control)
2DG 10 mM
azide 1 mM
HES7prom-luciferase activity
cross correlation
time (min)


Sup. Fig. 10

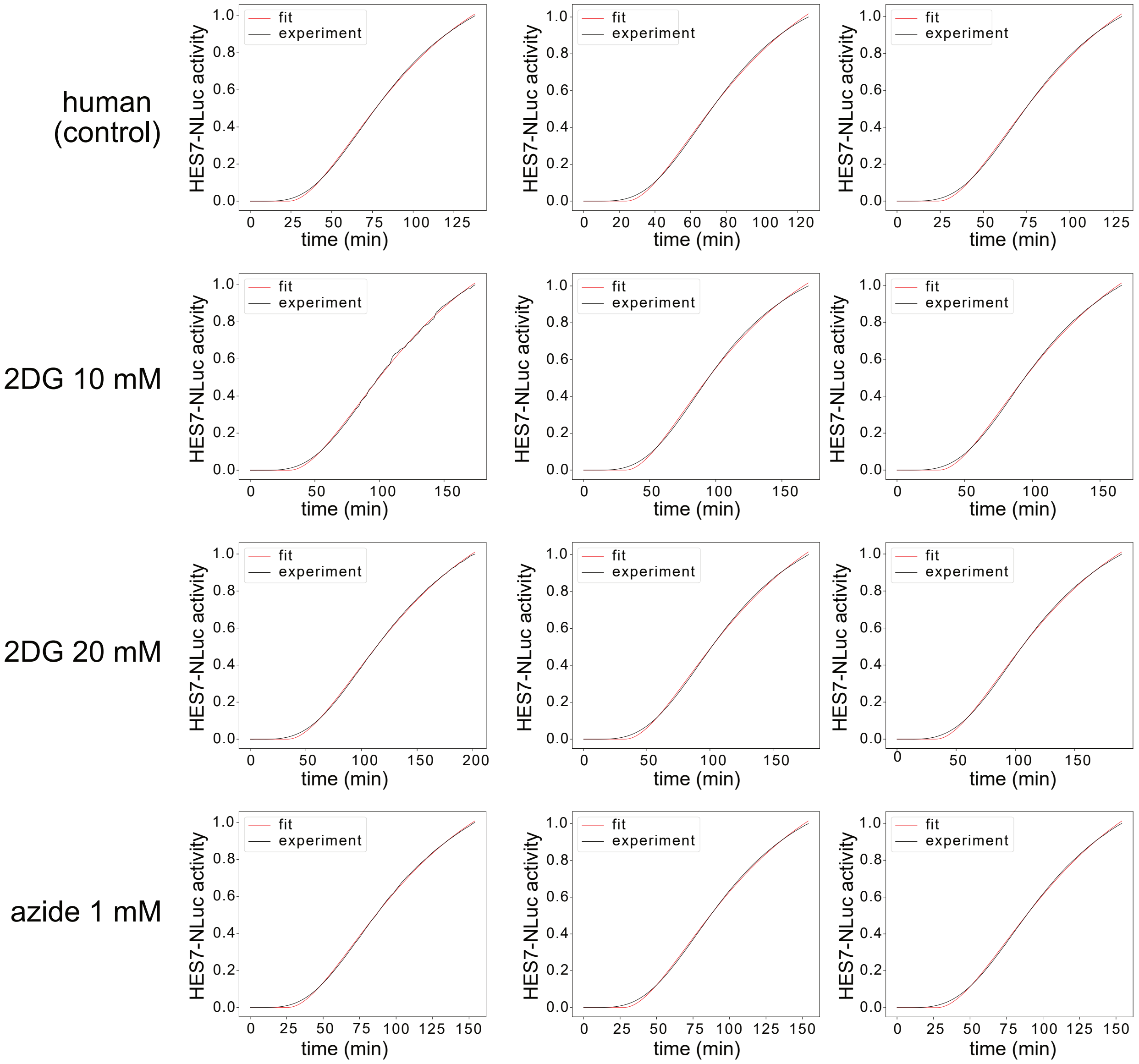
human (control)
2DG 10 mM
2DG 20 mM
azide 1 mM
fit
experiment
HES7-NLuc activity
time (min)

Sup. Fig. 11

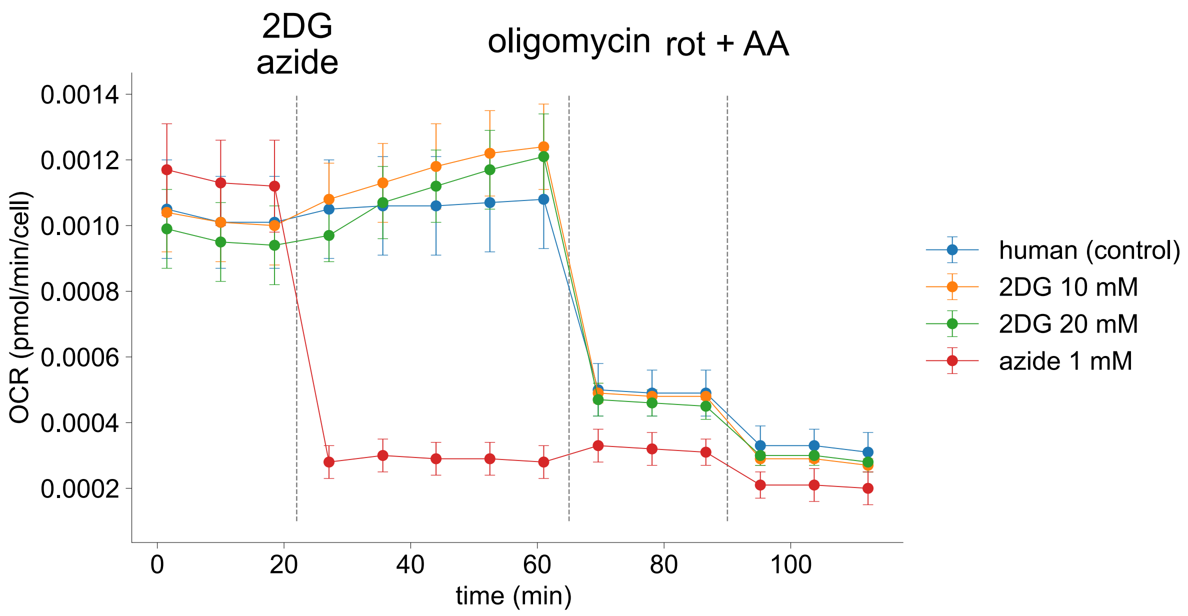
2DG
azide
oligomycin
rot + AA
0.0014
0.0012
0.0010
0.0008
0.0006
0.0004
0.0002
OCR (pmol/min/cell)
0
20
40
60
80
100
time (min)
human (control)
2DG 10 mM
2DG 20 mM
azide 1 mM

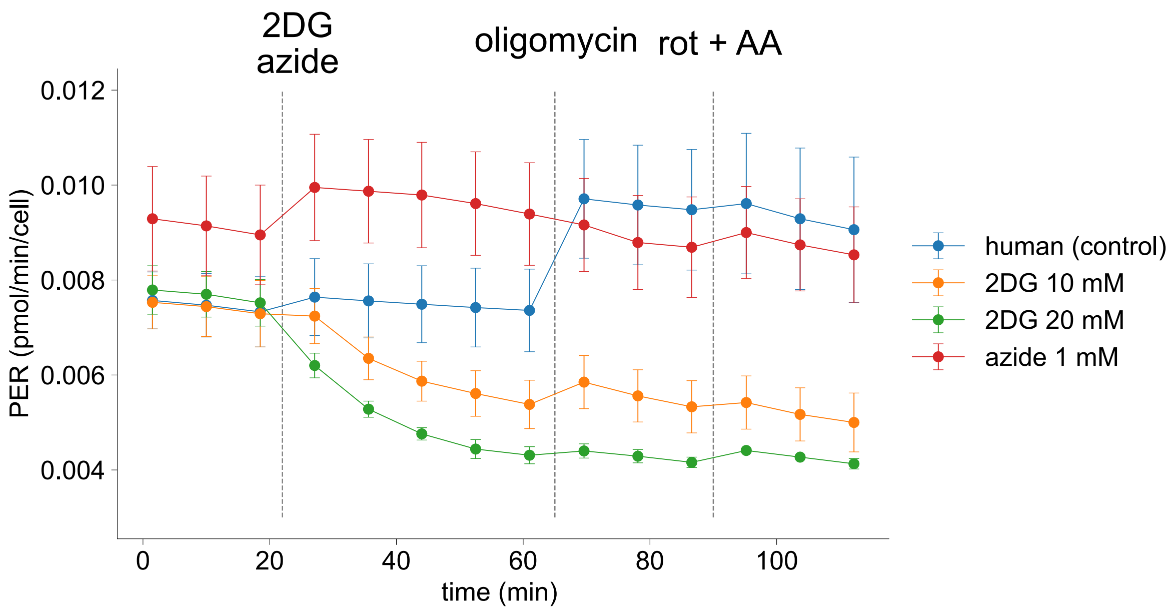
2DG
azide
oligomycin
rot + AA
0.012
0.010
0.008
0.006
0.004
PER (pmol/min/cell)
0
20
40
60
80
100
time (min)
human (control)
2DG 10 mM
2DG 20 mM
azide 1 mM

Sup. Fig. 12

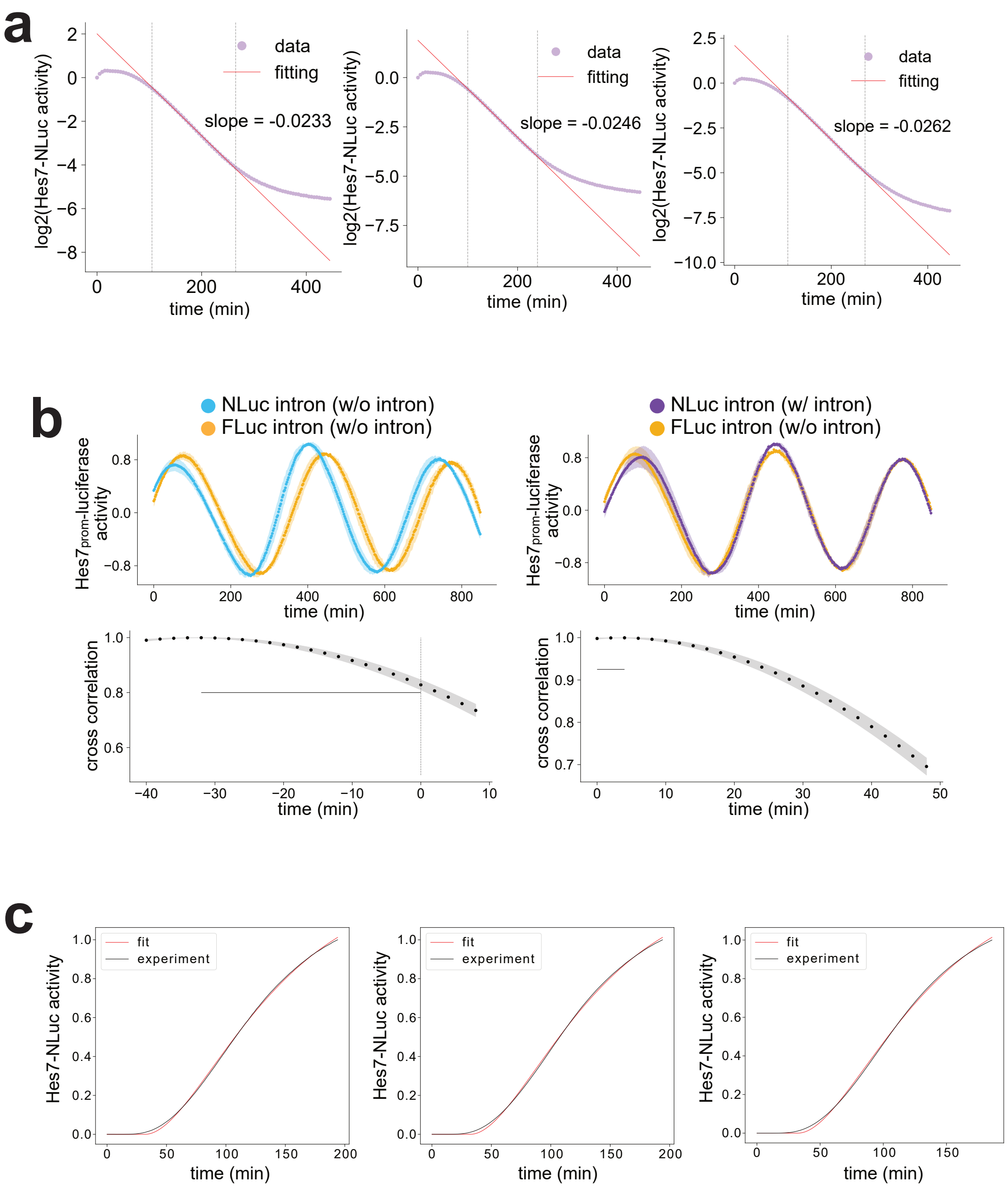
a
log2(Hes7-NLuc activity)
data
fitting
slope = -0.0233
time (min)
slope = -0.0246
slope = -0.0262
b
NLuc intron (w/o intron)
FLuc intron (w/o intron)
NLuc intron (w/ intron)
Hes7prom-luciferase activity
cross correlation
c
fit
experiment
Hes7-NLuc activity

Sup. Fig. 13
